# Enantiomeric Hydrogel-Manipulated Mechanotransduction Triggers Neurogenesis and Immunomodulation for Spinal Cord Repair

**DOI:** 10.1101/2025.01.10.632462

**Authors:** Ya Li, Yulin Wang, Xinyu Fang, Hengtong Zhang, Ruicheng Li, Rupeng Cai, Xiaohui Li, Zhaopu Yang, Xiaoyun Li, Zhongbing Huang, Guangfu Yin, Aijun Wang, Qiang Ao

## Abstract

Microenvironmental mechanics regulate morphogenesis and post-injury inflammation, however, the fragile mechanical strength and oxidative physiological environment hinder precise and consistent mechanical management after spinal cord injury (SCI). Here, we engineered self-assembling hydrogels of enantiomeric peptides with neural tissue-matching mechanical properties to persistently manipulate mechanosensing and mechanotransduction through stereo conformational recognition and consequent protein affinity difference. While hindering proliferation and morphogenesis in non-neural cells, D-hydrogel-induced intracellular tension relaxation triggered neurogenesis and ECM remolding in astrocytes, while simultaneously suppressing pro-inflammation and promoting pro-regeneration in microglia, which together enable neuroprotection from degeneration and enhance functional recovery in severe SCI rat models. These effects are mediated through neurogenic morphology changes resulting from cytoskeletal tension relaxation, leading to the opening of mechanosensitive ion channels in the cellular membrane, chromatin unfolding, and YAP nuclear translocation. This exclusive D-hydrogel-dependent neurogenesis, triggered by intracellular tension relaxation, revealed a neural-specific response to mechanical cues and provided a targeted tissue repair strategy for nerve injury.

**Teaser:** Intracellular tension relaxion activates morphogenesis specifically in neural cells through reversing neurogenic cellular morphology.

## Introduction

The absent neurogenesis and inflammation-driven secondary damage present two primary challenges in spinal cord injury (SCI) *(1, 2)*, resulting from highly specialized neural cytoskeleton precisely structured for efficient information transmission in the nervous system *(3)*. Over recent decades, efforts have focused on biochemical interventions, such as growth factor cocktail *(4)*, stem cells *(5, 6)* and diverse bioactive components *(7, 8)*, to regulate responsive neural cells and inflammation. However, the role of mechanical cues from extracellular matrix (ECM), where cells reside and are shaped, has been underemphasized. Mechanical signals from microenvironment regulate cell fate and shape morphogenesis *(9, 10)*, both of which are crucial for neurogenesis *(11, 12)* and immunomodulation *(13, 14)*. SCI dramatically alters mechanical properties as cysts, cavities, and glial scars form in the lesion *(15, 16)*. Nonetheless, our understanding of how mechanobiology influences neural cells and immune responses post-SCI remains limited. The major challenge is that traditional methods for engineering ECM mechanics, such as adjusting crosslinking density or adhesion ligand concentration *(17, 18)*, struggle to maintain a persistent mechanical gradient in the oxidative, enzyme-rich physiological environment particularly, within the weak strength range of central neural tissue (100 - 1000 Pa) *(19)*.

Cells detect microenvironment mechanics through recognition between adhesive ligands and membrane receptors *(20)*. Consequently, engineered enantiomeric ECM, as a stereoisomeric system that differentiates protein affinity *(21)*, enables refined and stable mechanosensing gradients while maintaining identical chemical and mechanical environments *(22)*, providing an advanced approach to exploring ECM mechanobiology. Compared to reported proliferation and differentiation improvement by L-enantiomeric ECM with stronger adhesion binding across multiple non-neural cell models *(23-27)*, our previous work reveals that D-enantiomeric ECM notably promotes several morphogenetic processes in neurons and glial cells *(28, 29)*, consistent with other reports on neural activity and differentiation enhancement *(30-32)*. This exclusive enantiomer-reversed mechanical response in neural cells strongly suggests that neurogenesis requires distinct mechanical properties for regeneration. Meanwhile, immune responses to ECM mechanics have also shown enantiomer-specific effects, with distinct levels of inflammation and even opposite regulatory effects *(33-36)*. However, studies utilizing different enantiomers in different immune cell models have yet to establish a consistent pattern. In central nervous system, microglia are specialized resident immune cells crucial in secondary inflammatory damage following SCI *(1, 37)*. Elucidating the immune effects of microglia responding to mechanical cues using enantiomeric ECM is also critical for defining the ECM mechanobiology essential for SCI restoration.

Here, we engineered enantiomeric hydrogels with mechanical properties matching neural tissues to manipulate mechanosensing and mechanotransduction through distinct protein affinity. D-enantiomeric hydrogel-induced intracellular tension relaxation triggered neurogenesis and ECM remolding in astrocytes, and simultaneously suppressed pro- inflammation while promoting pro-regeneration in microglia. In severe SCI rat models, D- hydrogel treatment significantly mitigated inflammatory damage and facilitated functional recovery. Single nucleus RNA sequence revealed that the neurogenesis and immunomodulation protected neuron from degeneration and enhanced synaptic reconstruction and neuronal communication. We explored the underlying mechanism of this exclusive D-hydrogel-dependent neurogenesis that neurogenic morphology changes resulting from cytoskeletal tension relaxation opens mechanosensitive ion channels in cellular membrane, unfolds chromatin, and translocates YAP into nuclei, revealing a neural-specific response to mechanical cues and providing targeted tissue repair strategy for nerve injury.

## Results

### Enantiomeric hydrogel preparation and characterization

To establish symmetrical chirality, we synthesized bioactive ligand-excluded tripeptide enantiomers, *N*-(9-fluorenylmethoxycarbonyl)-protected phenylalanine-phenylalanine- aspartic acid (Fmoc-^D^F^D^F^D^D and Fmoc-^L^F^L^F^L^D, Fig. S1), to self-assemble into supramolecular hydrogels (D-gel and L-gel, Fig. 1A, Fig S2A and B). The polarized light absorption signals with equal intensity but opposite signs in circular dichroism (CD) spectra (Fig. 1B) validated the mirror-symmetrical chirality between enantiomeric molecules and their supramolecular structures. The D-gel and L-gel have identical mechanical strength (G’ ∼ 800 Pa, G” ∼ 130 Pa, Fig. 1C), and stress relaxation profile (Fig. 1D), matching central nervous tissues *(19)*. Their rheological strength exhibited improvement in response to cerebrospinal fluid under oscillatory shear load (Fig. S2C), facilitating self-healing and injectability. Electron microscopy observations (Fig. 1E, Fig S2D and E) revealed the helical nanofiber assemblies in hydrogels with equivalent diameters and helical pitches (Fig. S2F and G) but opposite helical directions.

**Fig. 1.**
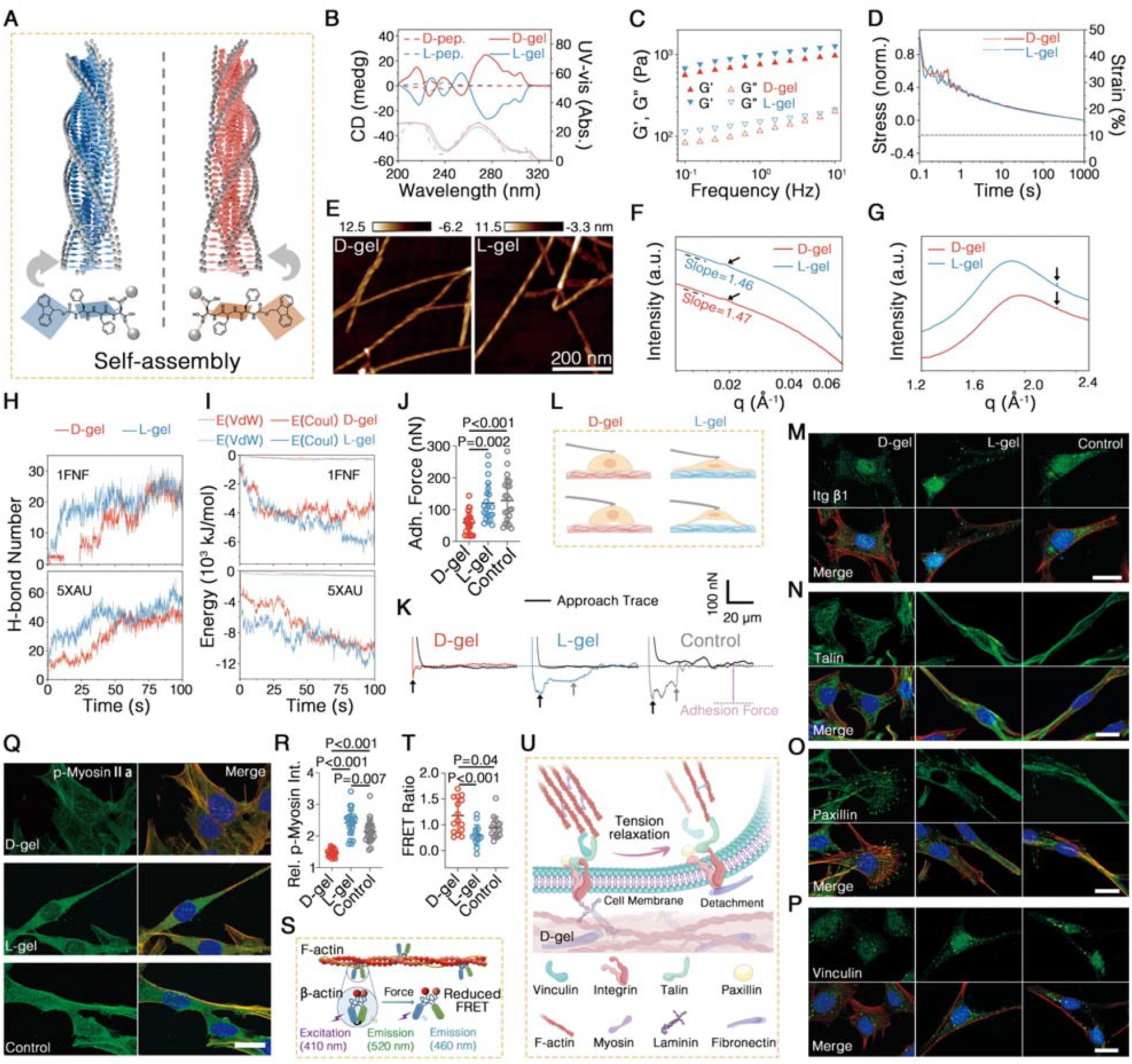
Engineered enantiomeric hydrogels manipulating intracellular tension. (**A**) Schematic representation of Fmoc-FFD enantiomers self-assembly into nanofiber hydrogels with mirror-symmetric helices. (**B**) CD spectra (in dark colors) and corresponding UV-vis spectra (in light colors) of Fmoc-FFD peptide enantiomers and corresponding hydrogels. (**C, D**) Rheological spectra of enantiomeric hydrogels presenting frequency-dependent strength (**C**) and stress relaxation (**D**). (**E**) AFM micrographs of enantiomeric hydrogels. (**F, G**) SAXS patterns (**F**) and WAXS patterns (**G**) of enantiomeric hydrogels. The scattering intensities were offset vertically for clarity; the characteristic peaks were indicated by black arrows. (**H, I**) Hydrogen bond formation (**H**) and interaction energy (**I**) between enantiomeric assemblies and fibronectin (1FNF) or laminin fragment (5XAU) from MD simulations. (**J, K, L**) Adhesion force magnitude (**J**) and force-probe displacement curves (**K**) between astrocytes and culture matrix measured by SCFS, with a schematic representation (**L**) of the experimental setup on enantiomeric hydrogels. (**M, N, O, P, Q, R**) Representative IF staining micrographs of astrocytes cultured on enantiomeric hydrogels, with marker in green, cytoskeleton in red, and nucleus in blue. Scale bar = 20 μm. Fluorescence intensity of p-Myosin was quantified in individual cells (**R**). (**S, T**) Schematic representation illustrating the process that cytoskeleton force reduces FRET efficiency (**S**), and FRET efficiency of astrocytes cultured on enantiomeric hydrogels (**T**). (**U**) Schematic representation illustrating D-gel-induced intracellular tension relaxation.

Furthermore, small-angle X-ray scattering (SAXS, Fig. 1F) confirmed analogous filament formation, exhibiting nearly identical Guinier region slopes (−1.46 and -1.47). The peak at 0.019 Å^-1^ (d-spacing 330 Å) suggested supramolecular hierarchical stacking and periodic arrangement, consistent with the helical structure. The identical peaks of enantiomeric wide-angle X-ray scattering (WAXS, Fig. 1G) at 1.9 Å^-1^ (d-spacing ∼ 3.3 Å) and 2.25 Å^-1^ (d-spacing ∼ 2.8 Å) corresponded to relaxed lateral stacking of β-strands and compactly ordered sequential stacking, respectively. The molecular dynamics (MD, Fig. S2H) simulation confirmed that the enantiomers assemble through π-π stacking of hydrophobic aromatic groups in the filament interior, while a few hydrogen bonds form between the hydrophilic ends of aspartic acid on the exterior. Collectively, we developed mirror-symmetrical enantiomeric hydrogels with identical non-stereochemical properties, central nervous tissue-equivalent mechanical strength, and cerebrospinal fluid-responsive self-healing.

### Enantiomer-dependent cellular adhesion mechanics

Since adhesion ligands were excluded in the FFD sequence, assembly-attracted proteins solely provide ECM anchors for mechanosensing *(38)*. We simulated the MD of physical interactions between enantiomeric assemblies and fibronectin/laminin, both of which possess integrin binding ligands *(39, 40)*, revealing that hydrogen bonds were predominant (Fig. S3) and formed fewer between D-assembly and both proteins (Fig. 1H). Interaction energy analyses corroborated that electrostatic energy (E_Coul_), contributing to hydrogen bonds *(41)*, dominated in the interactions, with Van der Waals energy (E_VdW_) nearly negligible (Fig. 1I). The negative sign and lower absolute E_Coul_ values validated the weaker attraction between D-assembly and both adhesive proteins. These simulations verify that D-gel possesses weaker protein affinity compared to L-gel, potentially facilitating detachment during cell adhesion.

We then quantified the adhesion force between cells and enantiomeric hydrogels using a single-cell force spectroscopy (SCFS). Standard cell culture plates treated for high adhesion were used as control. The adhesion force on D-gel was significantly reduced— 0.48-fold and 0.45-fold lower than that of the L-gel and control groups, respectively (Fig. 1J). The adhesion force distribution also differed (Fig. 1K), with the D-gel group exhibiting a single sharp peak corresponding to the central cell body, whereas the L-gel and control groups displayed an additional, broader peak attributed to the long protrusion characteristic of neural cells (Fig. 1L). These results substantiate that the cellular mechanosensing on engineered hydrogels is enantiomer-dependent. Adhesive anchors on D-gel are diminished and confined to the cell body, whereas on the L-gel group, adhesion is stronger and extends to the cell protrusions, resembling high-adhesion culture.

### Chirality mechanics-driven FA distribution and intracellular tension

Adhesion force is generated via focal adhesion (FA) *(42)*, within which transmembrane integrins sense and clutch ECM binding sites *(43)*. The immunofluorescence (IF) images (Fig. 1M) revealed a diffuse distribution of integrin β1 in astrocytes on D-gel with a spread cytoskeletal morphology and retracted protrusions, compared to the conspicuously clustering of integrin β1 at both the cell body and slender protrusions in the L-gel and control groups. Talin, linking integrins to cytoskeleton, also exhibited a diffuse distribution on D-gel whereas clustered orientation along protrusions on L-gel and high-adhesion surface (Fig. 1N). The lack of integrin and talin clustering on D-gel indicates the immaturity of FAs, resulting in unstable mechanical anchoring *(43)*. Paxillin (Fig. 1O), an FA scaffold protein that facilitates FA stabilization and indicates cytoskeletal movement *(44)*, formed clusters distributed along cytoskeletal edge without orientation on D-gel, in contrast to its alignment along the protrusions in the L-gel and control groups. Vinculin (Fig. 1P), a cytoskeleton linker in FAs *(45)*, exhibited loose perinuclear clustering on D-gel, while extending to the anterior protrusion with high clustering in the L-gel and control groups. These results suggest that, in astrocytes cultured on D-gel, immature and fragile FAs form, which are prone to rapid disassembly, providing unstable anchorage and disrupting cytoskeletal alignment. In contrast, mature, stable FA clusters form in the bipolar or tripolar morphology and neurogenic cellular protrusions of astrocytes cultured on L-gel and high-adhesion control surfaces. Additionally, this FA distribution pattern and cellular morphology on D-gel correspond to the SCFS characteristics of weaker adhesion force and the absence of a secondary protrusion force peak (Fig. 1L).

The mechanical force sensed by FAs is subsequently transmitted to cytoskeleton (*46*). The lowest level of p-myosin IIa (force generator in cytoskeleton *(47)*) on D-gel (Fig. 1Q and R) indicated cytoskeleton tension diminution. We then measured cytoskeleton force directly using Förster resonance energy transfer (FRET) technology *(48)*, where force-induced separation of the two fluorophores reduces the energy transfer efficiency (Fig. 1S). The increased FRET efficiency on D-gel (Fig. 1T, Fig. S4, 1.21-fold of the L-gel group and 1.11-fold of the control group) validated the significant intracellular tension relaxation. In summary (as illustrated in Fig. 1U), insufficient protein affinity results in the desorption of integrin-binding proteins from D-gel (Fig. 1H and I, Fig. S3) under cellular adhesive force (Fig. 1J-L), impairing FA maturation and stabilization (Fig. 1M-P) and leading to intracellular tension relaxation (Fig. 1Q-T, Fig. S4), compared to the high-adhesive L-gel. Although observed in astrocytes, this theory, based on enantiomer-dependent protein affinity, can be extended to other cell types. We successfully engineered mechanosensing and mechanotransduction manipulation using enantiomeric hydrogels.

### Intracellular tension relaxation facilitates morphogenesis in neural cells

Subsequently, we assessed the impacts of manipulated mechanotransduction on morphogenesis processes in astrocytes. Consistent with our previous findings *(29)*, astrocytes cultured on D-gel with low intracellular tension exhibited significantly enhanced viability (Fig. 2A, Fig. S5A), proliferation (Fig. 2B and C, Fig. S5B and C), and survival (anti-apoptosis, Fig. 2D, Fig. S5D), compared to the L-gel group with high tension. Moreover, the increased mitochondrial membrane potential (MMP, Fig. 2E, Fig. S5E) and elevated adenosine triphosphate (ATP) level (Fig. 2F) indicated that intracellular tension relaxation protected mitochondria from depolarization and facilitated energy metabolism in astrocytes. Cellular migration was also accelerated (Fig. S5F) with a rate of ∼31% after 24 h, 1.7-fold of L-gel group (Fig. 2G). Intracellular tension relaxation comprehensively promotes fundamental morphogenesis processes in astrocytes.

**Fig. 2.**
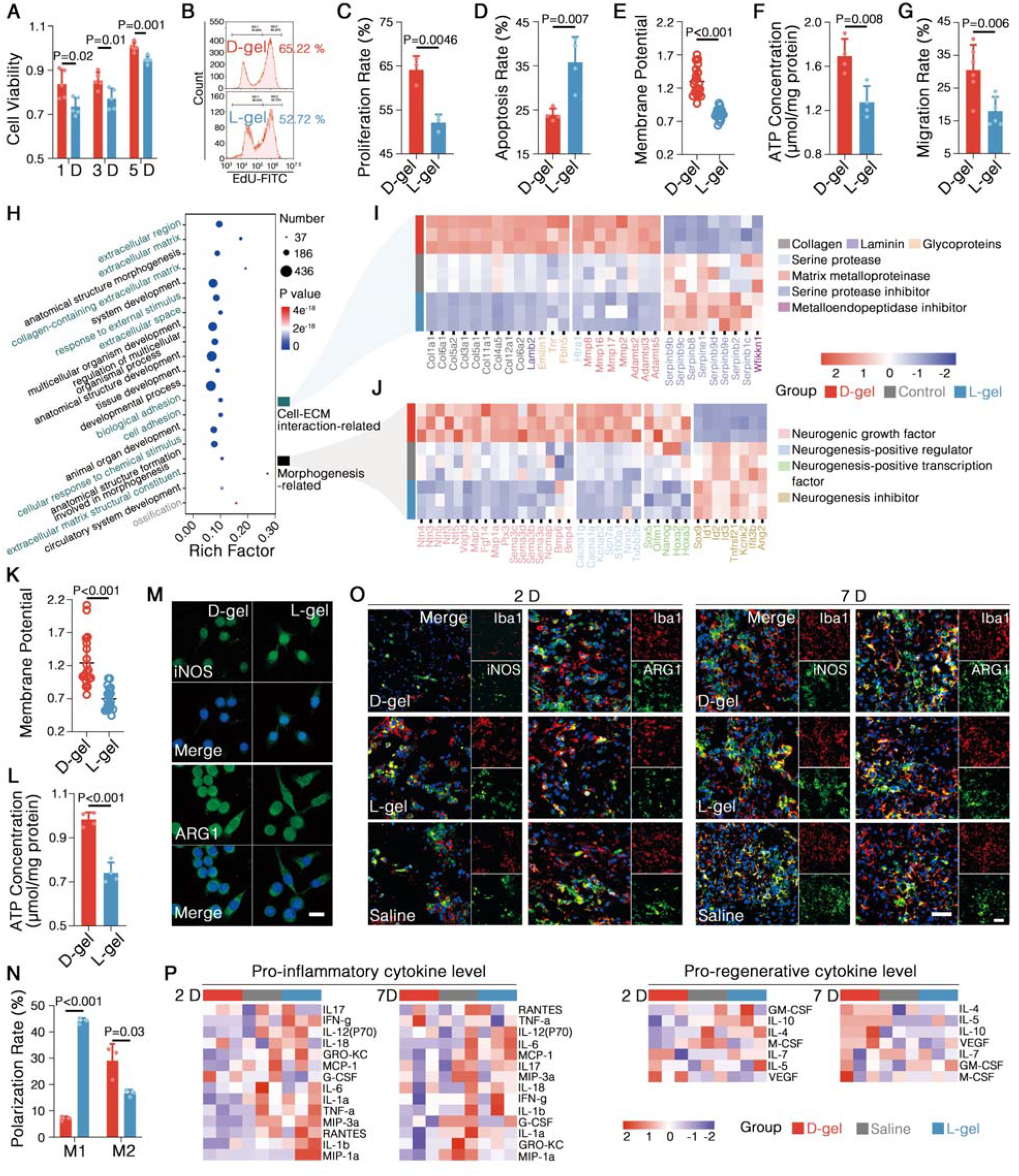
Intracellular tension relaxation triggers morphogenesis in astrocytes and immunomodulation in microglia. (**A**-**G**) Morphogenesis process of astrocytes cultured on enantiomeric hydrogels: cellular viability (**A**), flow cytometry histogram of proliferating cells (**B**), proliferation rate (**C**), apoptosis rate (**D**), mitochondrial membrane potential (**E**), ATP concentration (**F**), and migration rate in 24 h (**G**). (**H, I, J**) Top 20 GO terms enriched in upregulated astrocyte genes in the D-gel group compared to the L-gel group, based on RNA-seq (**H**), with corresponding heatmaps showing gene expression related to Cell-ECM interactions (**I**) and morphogenesis (**J**). (**K, L**) Mitochondrial membrane potential (**K**) and ATP concentration (**L**) of microglia cultured on enantiomeric hydrogels. (**M**) Representative IF staining micrographics of microglia. iNOS labels M1 pro- inflammatory polarization, Arg1 marks M2 pro-regenerative polarization. DAPI- stained nucleus is in blue. Scale bar = 20 μm. (**N**) Polarization rate of microglia cultured on enantiomeric hydrogels based on flow cytometry assay. (**O**) Representative IF staining micrographics of spinal cord lesion on day 2 and 7 postoperatively, Iba1 labels microglia and DAPI-stained nucleus is in blue. Scale bar = 50 μm. (**P, Q**) Heatmaps of pro-inflammatory (**P**) and pro-regenerative (**Q**) immune cytokine levels in spinal cord lesion at day 2 and 7 postoperatively.

To gain a deeper insight, we analyzed global gene expression profile of astrocytes under manipulated intracellular tension (high-adhesion culture plate was again used as high-tension control here, Fig. S6A and B). The transcriptome on D-gel underwent substantial changes compared to the similar profiles of the L-gel and control groups (Fig. S6C). The upregulated genes were annotated (Fig. S6D) and analyzed through gene ontology (GO) enrichment (Fig. 2H), revealing their predominant involvement in cell-ECM interactions (9 of the top 20 terms) and morphogenesis (10 of the top 20 terms). Further clustering of cell-ECM interaction-related genes (Fig. 2I) showed that D-gel upregulated genes encoding ECM components and ECM proteases, while downregulating protease inhibitors. Meanwhile, clustering of morphogenesis-related genes (Fig. 2J) revealed that D-gel upregulated genes encoding growth factors, neurogenesis-positive regulators and transcription factors, while downregulating neurogenesis inhibitors. These results highlight that intracellular tension relaxation initiates a dynamic process of extracellular matrix construction and remodeling, while triggering neurogenesis, which is essential for nerve regeneration.

### Intracellular tension relaxation induces immunomodulation in microglia

To substantiate the mechanobiological manipulation of microglia by enantiomeric hydrogels, we first assessed cytoskeletal tension. Similar to astrocytes, microglia cultured on D-gel exhibited increased FRET efficiency (Fig. S7), corroborating lower intracellular tension and confirming that D-gel induces tension relaxation indiscriminately across all cell types through stereo conformational misrecognition and consequent low protein affinity. Regarding immune response, most genesis processes of microglia showed no significant nor coherent difference on enantiomeric hydrogels (Fig. S8). However, the apoptosis on D-gel is decreased (Fig. S9) and MMP and ATP levels are significantly increased (Fig. 2K and L, Fig. S10), hinting that microglia are activated by enantiomeric gels. IF staining showed that D-gel downregulated the pro-inflammatory subtype marker inducible nitric oxide synthase (iNOS) and upregulated the pro-regenerative subtype marker arginase (Arg, Fig. 2M). Flow cytometry revealed that the pro-inflammatory subtype accounted for 6.98% in the D-gel group, 0.16-fold lower than that in the L-gel group, while the pro-regenerative subtype accounted for 29.04%, 1.74-fold higher than in the L-gel group (Fig. 2N, Fig. S11). These in vitro results suggest that intracellular tension relaxation inhibits pro-inflammatory polarization and promotes pro-regenerative polarization in microglia.

To evaluate *in vivo* responses, enantiomeric hydrogels were injected into severe rat SCI models and gelled *in situ* at the lesion core. IF staining of the lesion core at acute phase (postoperative day 2) and subacute phase (day 7) exhibited the lower iNOS labelling and higher Arg labeling after D-gel treatment compared to the L-gel and saline groups, further corroborating the intracellular tension relaxation induced pro-inflammatory polarization inhibition and pro-regenerative polarization promotion in microglia (Fig. 2O). The immune cytokine levels of spinal cord lesion were analyzed using high-throughput protein microarray technology (Fig. 2P). After D-gel injection, the pro-inflammatory cytokines were massively downregulated during both acute phase and subacute phase. Meanwhile, pro-regenerative cytokines were generally upregulated during the subacute phase, with no significant differences observed in the acute phase, which aligns with the inflammation timeline, where regenerative responses occur later. Collectively, these results demonstrate that D-gel-induced intracellular tension relaxation suppresses post-injury inflammation and promotes pro-regenerative polarization along with the corresponding cytokine secretion, preventing secondary inflammation damage following SCI.

### SCI repair under enantiomer-engineered ECM mechanotransduction

The long-term rehabilitation of severe SCI after enantiomeric hydrogel injection (28 days) was investigated subsequently. In the D-gel group, Basso-Beattie-Bresnahan (BBB) scores, as well as movement velocity and activity time in the open field (Fig. 3A-D) were significantly increased, compared to the L-gel and saline groups. Meanwhile, the response latency to thermal stimulus (Fig. 3E) was significantly reduced. The footprints exhibited restored movement with spread toes and sustained coordination of forelimbs and hindlimbs (Fig. 3F, Fig. S12B). Additionally, the electrophysiological transduction (Fig. 3G) was successfully reconstructed with enhanced amplitude and diminished latency in the D-gel group (Fig. 3H and I), while no signals were observed in the saline group. These results demonstrate that D-gel-mediated intracellular tension relaxation facilitates locomotor recovery and accelerates sensory feedback after SCI.

**Fig. 3.**
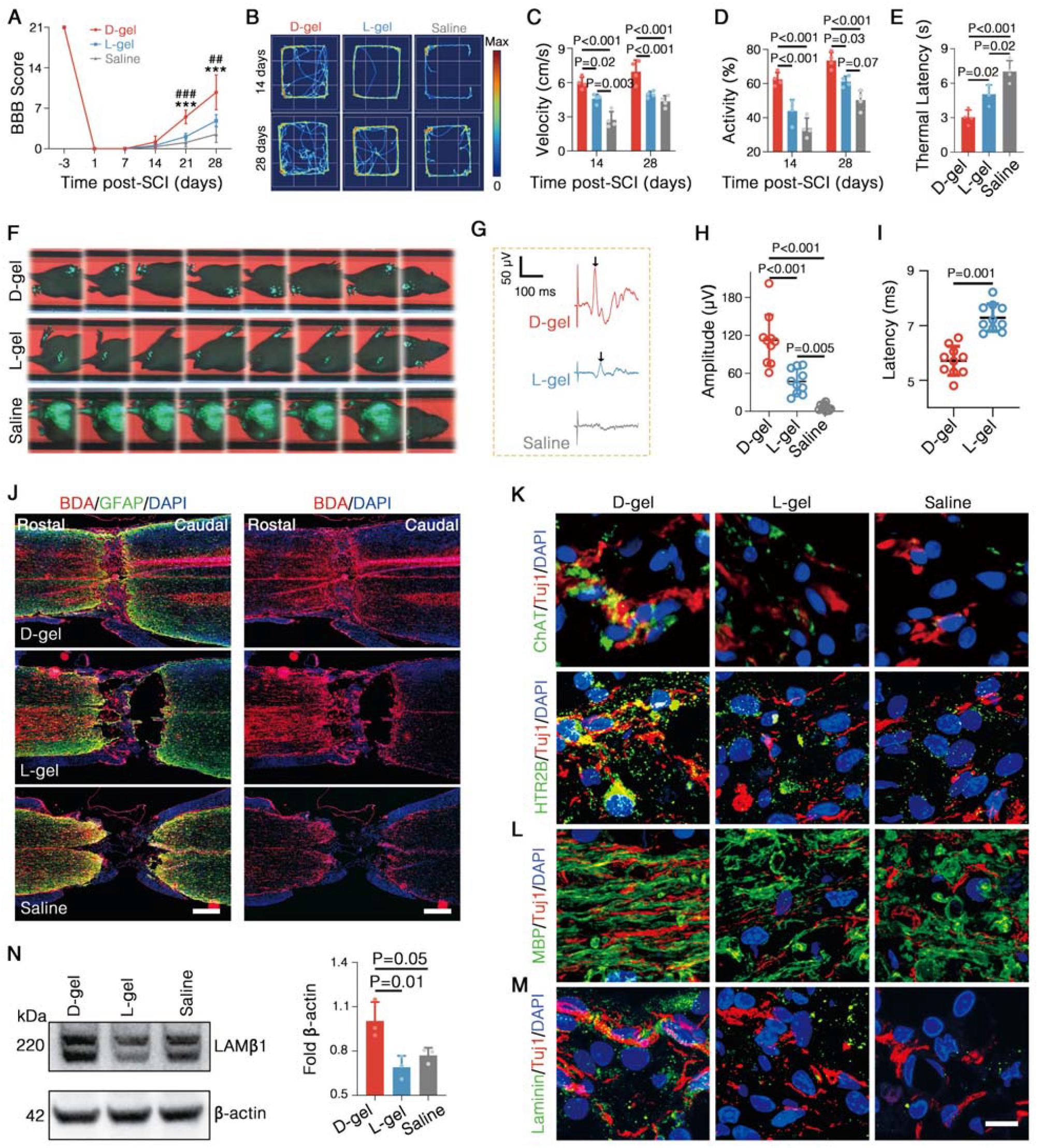
SCI repair under enantiomer-manipulated mechanotransduction cues. (**A**-**D**) Locomotion recovery after SCI: BBB scores (##*P<0*.*01*, ###*P<0*.*001*, compared with L-gel group; \*\*\**P<0*.*001*, compared with Saline group) (**A**), activity track heatmap (**B**) and statistical analyses of movement velocity (**C**) and activity time percentage (**D**) in the open field. (**E**) Latency response to thermal stimulus at day 28. (**F**) Representative motion trajectories of rat models at day 28. (**G**-**I**) Motor evoked potentials (**G**) and statistical analyses of amplitude (**H**) and latency (**I**) at day 28. (**J**) Fluorescent micrographics of longitudinal spinal cord sections. GFAP indicates astrocytes (green), BDA indicates labeled axons (red), and DAPI indicates nuclei (blue). Scale bar = 500 μm. (**K, L, M**) Representative IF staining micrographics of spinal cord. Tuj1 labels neuron and DAPI labels nuclei. Scale bar = 10 μm. (**N**) WB results and statistics of laminin in injured spinal cord.

Biotinylated dextran amine (BDA) was administered to anterogradely trace the axon regrowth. In rat injected with saline or L-gel, clear cavitation, resulting from secondary inflammation damage, and few regrown axons was observed within the lesion, whereas we observed substantial tissue and some regrown axons across the lesion core in the D-gel group (Fig. 3J, Fig. S13A). The IF staining exhibited that the lesion core tissue was primarily constituted of neurons with a lack of astrocytes (Fig. S13B). And no apparent astrocyte scars formed around the lesion in the D-gel group, compared to the dense population of reactive astrocytes at the lesion borders in the L-gel and saline groups, which strongly inhibits axonal regeneration *(49)*. Both cholinergic neurons (ChAT) and serotonergic neurons (5HT), locomotor function-related neurons, in the lesion were also observed improved in the D-gel group (Fig. 3K, Fig. S14). The demyelination of white matter in caudal lesion was determined according to myelin basic protein (MBP, Fig. 3L, Fig. S15). MBP exhibited alignment and wrapping around axons in the D-gel group, whereas disordered and scattered in the L-gel and saline groups, revealing effective anti- demyelination after D-gel injection. Additionally, ECM component laminin was significantly increased and distributed around nerves in the spinal cord (Fig. 3M and N, Fig. S16), indicating neuron supporting of the reconstructed ECM. Our observations suggested that D-gel treatment enabling intracellular tension relaxation significantly enhances spinal cord recovery, inhibiting glial scar formation and demyelination.

### Single-nucleus transcriptome atlas following SCI

To obtain a deeper insight into the single-cell characteristics following SCI under the enantiomer-engineered ECM mechanical stimulus, we performed single-nucleus RNA-sequence (snRNA-seq) on spinal cords collected at day 10 post-injury. Coarse clustering identified 10 cell types (Fig. S17), among which astrocytes, microglia, and neurons were further refined with hierarchical clustering due to their dramatic changes after injury (*49*). We identified 8 astrocyte clusters using unbiased clustering (Fig. 4A) and analyzed their gene characteristics through KEGG pathway enrichment. Cluster 1-3 were diverse reactive astrocytes as we discussed in Supplementary Text (Fig. S18) and showed little fluctuation in numbers across three groups. Cluster 6-8 were excluded here due to their minimal cell numbers and lack of significant changes. In cluster 4, substantially upregulated neurodegenerative disease-related pathways and marker genes indicated pathological degeneration and metabolic dysfunction (Fig. 4B and C). Gene set variation analysis (GSVA) across three groups (Fig. 4D) showed downregulation of all neurodegenerative pathways in the D-gel group, which, along with the greatly reduced cluster 4 percentage (Fig. 4A), suggested that intracellular tension relaxation protected astrocytes from degeneration after SCI. In cluster 5, top upregulated morphogenesis-related pathways (Fig. 4E) strongly indicated their neurogenesis function, along with specific marker genes involved in neurodevelopment, growth factors, transcriptional factors, ECM components, and ECM proteases (Fig. 4F, Fig. S19). This gene expression characteristics of cluster 5 highly matches the astrocyte gene expression profile observed *in vitro* culture on D-gel through RNA-seq (Fig. 2H-J), both showing strong ECM remolding and neurogenesis activity. The percentage of cluster 5 in the D-gel group increased dramatically, reaching 4.43-fold of the L-gel group. Meanwhile, these morphogenesis-related pathways and genes were significantly upregulated in the D-gel group (Fig. 4G and H). These results provide single-cell evidence that intracellular tension relaxation effectively activates ECM remodeling and neurogenic activity in astrocytes following SCI.

**Fig. 4.**
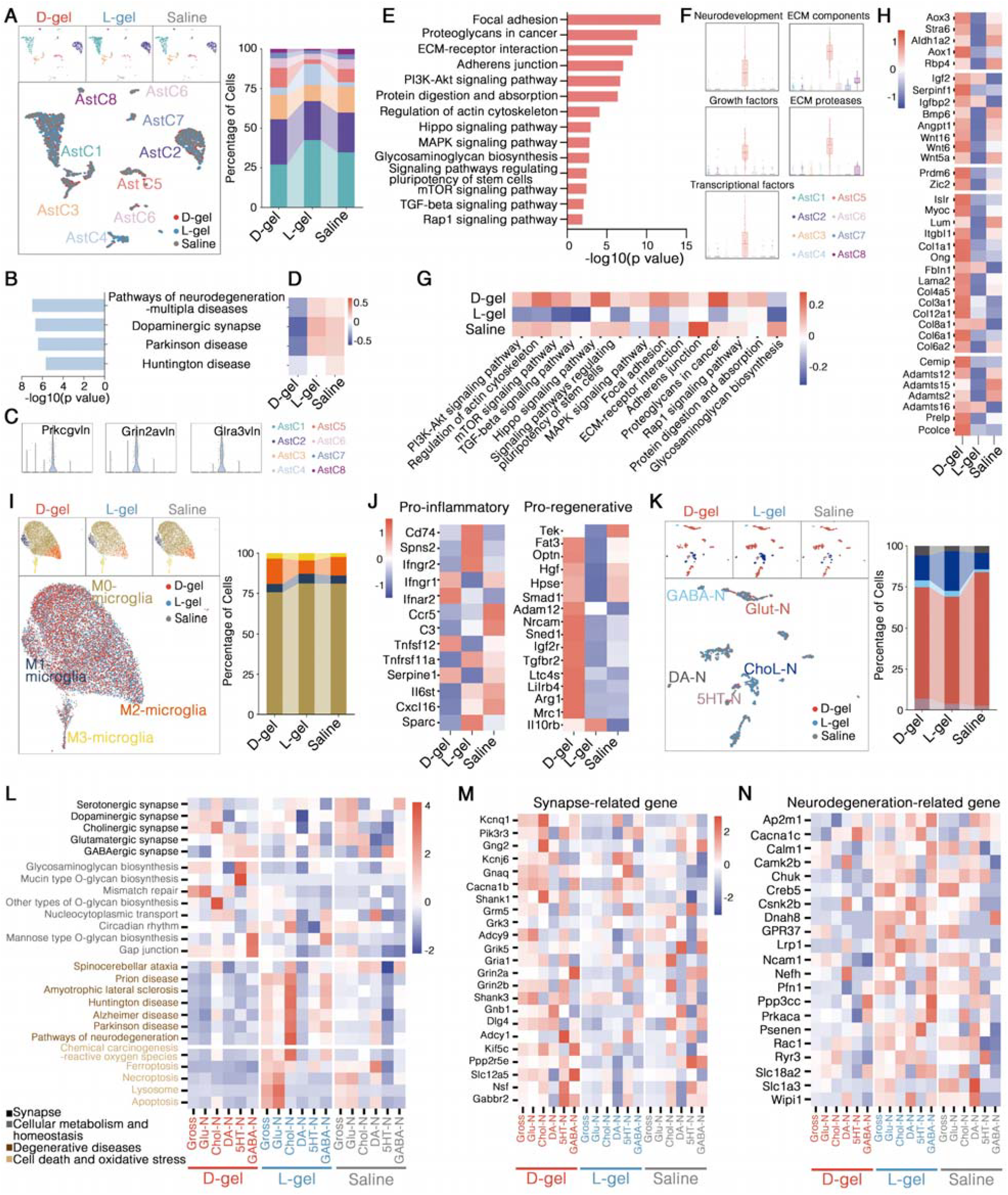
The snRNA-seq atlas of spinal cord at day 10 post-injury. (**A**) UMAP visualizations and proportions of astrocyte clusters. (**B, C, D**) Degeneration-related KEGG pathways (**B**) top enriched in AstC4, corresponding marker gene levels across different astrocyte clusters (**C**), and corresponding GSVA heatmap depicting the KEGG pathway levels in AstC4 across different groups (**D**). (**E, F, G, H**) Neurogenesis-related KEGG pathways (**E**) top enriched in AstC5, corresponding GSVA of marker genes classified according to gene function across different astrocyte clusters (**F**), and corresponding GSVA heatmap (**G**) depicting the KEGG pathway levels in AstC5 across different groups. Panel (**H**) is corresponding morphogenesis-related gene level heatmap across different groups. (**I, J**) UMAP visualizations and proportions of microglia clusters (**I**) and immune gene level heatmap across different groups (**J**). (**K, L, M, N**) UMAP visualizations and proportions of neuron clusters (**K**), GSVA heatmap of top enriched KEGG pathways characteristic in all neuron clusters (**L**), synapse-related gene level (**M**) and neurodegeneration-related gene level (**N**) heatmaps of each cluster and neuron gross across different groups.

Microglia were clustered as M0-activated, M1-proinflamatory, M2-proregenerative, and M3-others, based on reported marker genes *(50)* (Fig. 4I, Fig. S20). M0 cells were most abundant, slightly decreased in the D-gel group. M1 showed minimal differences, while M2 significantly increased in the D-gel group, 1.9 times higher than L-gel. Meanwhile, the pro-inflammatory genes were generally downregulated and the pro-regenerative genes were massively upregulated in the D-gel group (Fig. 4J). These results further validated the anti-inflammatory and pro-regenerative responses of microglia to D-gel-mediated intracellular tension relaxation, confirming to the *in vitro* and *in vivo* results. Neurons were clustered as glutamatergic neuron (Glu-N), GABAergic neuron (GABA-N), cholinergic motor neuron (Chol-N), dopaminergic neuron (DA-N), and serotonergic neuron (5HT-N), according to their transmitters (Fig. 4K) *(51)*. We classified specific KEGG pathways in each cluster and compared their gene set levels across three groups and corresponding clusters through GSVA (Fig. 4L-N, Fig. S21), revealing that D-gel injection upregulated pathways and genes involved in synapse, cellular metabolism and homeostasis, and massively downregulated pathways and genes involved in degenerative diseases, cell death and oxidative stress. These results further corroborate that intracellular tension relaxation induced by D-gel provides neuroprotection and mitigates degeneration after SCI.

### Single-nucleus transcriptome atlas: cellular communication

To determine how these cells interact, we performed CellChat to analyze the cellular communication patterns among these neurons, astrocyte cluster 5 (AstC5), M1 and M2 microglia. The overall interaction intensity network of the ligand-receptors (Fig. 5A) highlighted the strongest interactions between neurons, with AstC5 and M1 and M2 microglia influencing neurons. Enantiomeric hydrogel-treatment groups exhibited comprehensive communication, which is higher in the D-gel group, whereas the saline group showed minimal interaction, with GABA-N, 5HT-N, and DA-N remaining isolated. The heatmap of significant ligand-receptor pairs (Fig. S22-24) and the corresponding outgoing/incoming signaling patterns (Fig. 5B, Fig. S25) reveal the signaling pathways where one cell type provides ligands and another provides receptors. Neurons interacted with each other through neurotransmitter/synapse (Glutamate, NRXN, ADGRL, GABA- B) and intercellular adhesion/signaling (PTPR, NEGR, NCAM). Further comparing the intensity of these pathways across all three groups (Fig. 5C, Fig. S26A) revealed that the D-gel group exhibited more ligand-cell to receptor cell pairs and stronger interaction intensities, represented by a greater number and thicker interaction lines in the figure, indicating enhanced synaptic signaling and inter-neuronal communication. This result was consistent with the previous observation that D-gel facilitated functional recovery and electrophysiological reconstruction and upregulated synapse-related genes (Fig.3A-I, 4M).

**Fig. 5.**
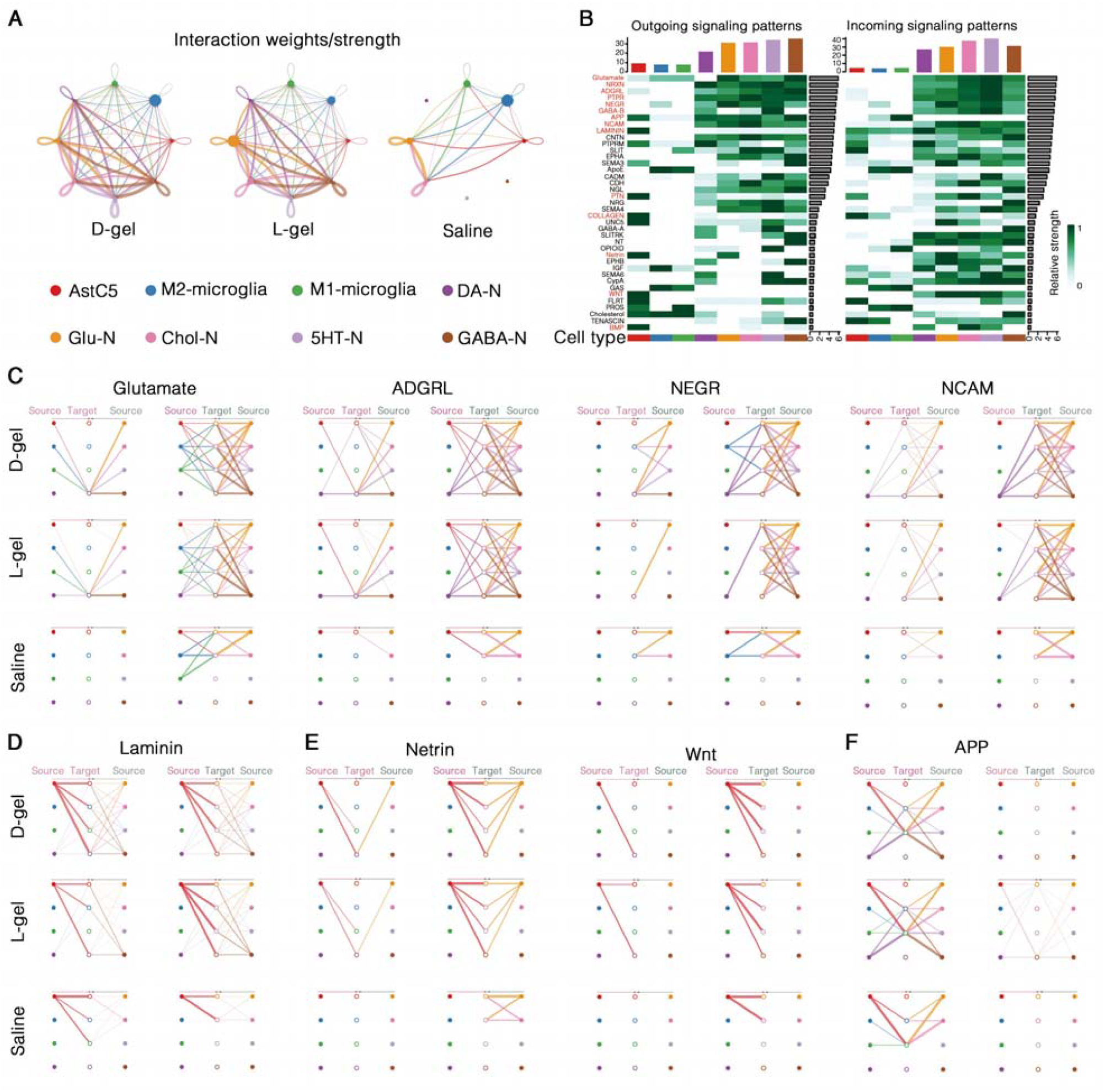
Cellular communication based on snRNA-seq atlas. (**A**) Gross interaction strength network between AstC5, M2-microglia, M1-microglia, DA-N, Glu-N, Chol-N, 5HT-N, and GABA-N. (**B**) High contributed signaling pathway patterns of the D-gel group. Outgoing signaling patterns indicate ligand-providing cell types, and incoming signaling patterns indicate receptor-providing cell types. (**C, D, E, F**) Hierarchical diagram of ligand-receptor interactions within signaling pathways: pathways between neurons (**C**), pathways through which AstC5 affects others (**D**), pathways through which AstC5 influences neurons exclusively (**E**), pathways through which other cells impact microglia (**F**). Solid circles represent ligand-providing cells, hollow circles represent receptor-providing cells, and thicker lines indicate stronger signals.

AstC5 exerted its effect on neurons and microglia through Laminin and Collagen pathways (Fig. 5D, Fig. S26B), with enhanced intensities in the D-gel group, where AstC5 served as the primary ligand provider and other cells provided the corresponding receptors. As described above, ECM components, particularly laminin, were significantly upregulated by D-gel both *in vitro* and *in vivo* (Fig. 2I, 3M and N). These findings collectively validate that ECM remolding induced by D-gel-mediated intracellular tension relaxation promotes nerve repair comprehensively. Additionally, AstC5 influenced neurons exclusively through PTN, Netrin, WNT, and BMP pathways (Fig.5E, Fig. S26C), where PTN and BMP are growth factors, Netrin is neural navigation cue, and WNT is a key developmental signaling pathway, all of which are crucial for neurogenesis. Their improvement in the D-gel group indicated that D-gel-upregulated factors related to growth and development (Fig. 2J, 4H, Fig. S21A) successfully communicated with neurons, demonstrating intracellular tension relaxation-maneuvered neurogenesis on a deeper level.

APP (amyloid precursor protein) pathway, closely associated with the pathogenesis of neurodegeneration diseases, was the primary pathway through which other cells affected microglia (Fig. 5F), indicating that the degeneration of astrocytes and neurons activates microglia, particularly inflammation (M1-microglia), which is decreased in the D-gel group compared to L-gel. Additionally, as we described above, D-gel downregulates the neurodegeneration disease-related genes and prevents astrocytes and neurons from degeneration (Fig. 4D and N). These findings collectively demonstrated that the D-gel further mitigates inflammation damage through neuroprotection and anti-degeneration, in addition to direct pro-inflammatory polarization inhibition and pro-regenerative polarization improvement.

Overall, *in vivo* experiments demonstrated that D-gel-mediated intracellular tension relaxation maneuvered ECM remolding and neurogenesis in astrocytes and facilitated synaptic signaling communication of neurons. The neurodegeneration after SCI was also inhibited, which mitigated inflammation response. Meanwhile, pro-inflammatory polarization and gene expression of microglia was suppressed and pro-regenerative polarization and gene expression was enhanced directly. These intracellular tension relaxation-driven effects collectively promote spinal cord repair post-injury.

### Neurogenic morphology changes maneuver neurogenesis through epigenetic remolding and mechanosensitive ion channel

However, it is generally accepted that mechanical morphogenesis is driven by strong adhesion-induced high intracellular tension *(52, 53)*, particularly in studies focusing on how L-chiral ECM with high adhesion promotes non-neural cell proliferation *(23, 25, 54)*, and that the cytoskeletal contractility transfers this tension to the nucleus and cell membrane, activating mechanosensitive pathways *(55, 56)*. To further validate the neurogenesis activation by intracellular tension relaxation, blebbistatin—a myosin II ATPase inhibitor disrupting actomyosin contractility—was solely used to relax cytoskeletal tension in astrocytes. The similarly increased cellular activity and proliferation (Fig. S27) after treatment directly, highlighted the exceptional distinct response of neural cells to mechanical signals. Considering the remarkable morphology transformation with disoriented spreading and retracted protrusion on D-gel, we hypothesized that the key factor was the variation of neurogenic morphology.

We reconstructed 3D models of the cell membrane and corresponding nucleus of astrocytes (Fig. 6A) to quantify their volumes. Statistical analyses (Fig. 6B) revealed a positive correlation between cell volume and nuclear volume and that both cell and nuclear volumes in the D-gel group significantly increased, reaching 1.74-fold and 2.29- fold of the L-gel group, respectively. These results indicated osmotic swelling of neural cells and their nuclei after intracellular tension relaxation, which could change cell membrane tension and nucleic stiffness *(57)*. We then used a live cell fluorescent probe (Flipper-TR^®^) combined with fluorescence lifetime imaging microscopy (FLIM) to assess the cell membrane tension. The significantly prolonged fluorescence lifetime on D-gel demonstrated the increased membrane tension (Fig. 6C and D), which typically leads to membrane rigidity enhancement and fluidity reduction. This was validated by significantly increased fluorescent half-recovery time after photobleaching (FRAP) on the cell membrane, (Fig. 6E and F), further confirming the elevated cell membrane tension, which can pull open mechanosensitive ion channels *(58, 59)*. IF staining and Western blot (WB) assays revealed the significant clustering and upregulated expression of Piezo 1 and transient receptor potential cation channel subfamily V member 4 (TRPV4, Fig. 6G and H), suggesting cooperatively enhanced ion channel efficiency and opening (*58*). The resulted Ca^2+^ influx (Fig. 6I and Fig. S28) was subsequently demonstrated improved, which can initiate morphogenesis through multiple Ca^2+^-dependent cascades *(60, 61)*. In the nucleus, the nuclear intermediate filament Lamin A/C, which carries nuclear tension and transfers it into chromatin condensation *(62)*, exhibited significant upregulation in the D-gel group (Fig. 6J and M), corroborating the improved nuclear stiffness and suggesting chromatin unfolding. The reduced histone deacetylase (HDAC, an enzyme catalyzing chromatin folding *(63)*, Fig. 6k and m) demonstrated the chromatin unfolding and enhanced transcriptional accessibility. The improved nuclear translocation of Yes-associated protein (YAP, a mechanosensitive transcriptional activator *(64)*, Fig. 6L and M) validated transcriptional activation, aligning with the substantial transcriptome variation (Fig. 2J).

**Fig. 6.**
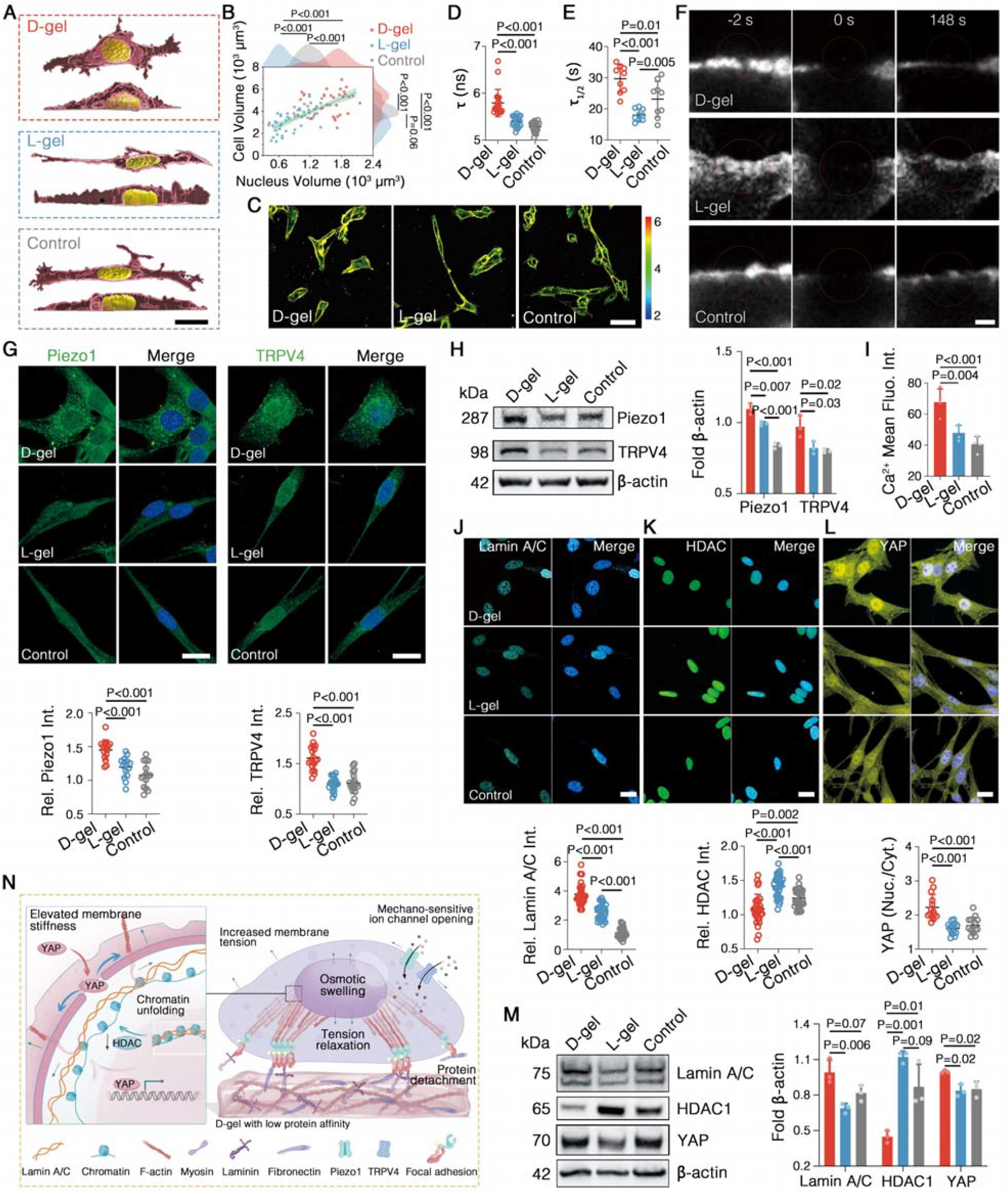
Neurogenic morphology variation and downstream mechanical signaling. (**A, B**) Computational reconstruction of astrocyte cell nuclei and cell membranes (**A**) and corresponding volume statistics (**B**). (**C, D**) FLIM heatmap of astrocytes staining with Flipper-TR^®^ (**C**) and corresponding fluorescence lifetime statistics (**D**). (**E, F**) Fluorescent half-recovery time after photobleaching (**E**) and representative micrographs (**F**). (**G**) Representative IF staining micrographs of astrocytes cultured on enantiomeric hydrogels with DAPI-stained nuclei in blue and corresponding quantification in individual cells. (**H**) WB results and statistics of mechanosensitive ion channels in astrocytes. (**I**) Fluorescence intensity statistics of stained Ca^2+^ in astrocytes. (**J, K, L**) Representative IF staining micrographs of astrocytes for Lamin A/C (**J**), HDAC (**K**), and YAP (**L**) with DAPI-stained nuclei in blue and corresponding quantification in individual cells. The ratio of nuclear intensity to cytoplasmic intensity of YAP is calculated. (**M**) WB results and statistics of Lamin A/C, HDAC, and YAP in astrocytes. (**N**) Schematic representation illustrating that intracellular tension relaxation opens mechanosensitive ion channel, unfolds chromatin, and activates transcription through increasing membrane tension and nuclear stiffness. Scale bars, 30 μm (**A**), 50 μm (**C**), 2 μm (**F**) 20 μm (**G, J, K, L**).

Collectively, as illustrated in Fig. 6N, D-enantiomeric hydrogel-driven intracellular tension relaxation disrupts polarized neurogenetic morphology, leading to osmotic swelling. The expanding membrane increases both cell membrane tension and nuclear stiffness, which opens mechanosensitive ion channels, unfolds chromatin, and triggers transcription, ultimately maneuvering neurogenesis.

## Discussion

Our work demonstrates that intracellular tension relaxation induced by ECM mechanics activates neurogenesis in astrocytes and modulates immune responses of microglia, leading to neuroprotection, synapse signaling enhancement, and inflammatory damage inhibition, consequently facilitating functional recovery from SCI. The neurogenesis activation in astrocytes aligns with our previous study *(29)*, where D-enantiomeric hydrogel enhances morphogenetic processes in multiple neural cells, including immortalized neuron, neuron-like cell, and Schwann cell, further validating that intracellular tension relaxation maneuvers neurogenesis, despite not observing this phenomenon directly in spinal cord neurons due to their advanced differentiation. Meanwhile, numerous studies have shown that non-neural cells typically exhibit inhibited morphogenesis, particularly in proliferation, when cultured on D-enantiomeric matrices *(23-27, 52-54)*. These findings imply that intracellular tension relaxation-induced morphogenesis is a neural-specific response, which is further highlighted by the mechanism that the consequent change of neurogenic morphology activates the downstream mechanical signaling. During neural cell maturation, as protrusions are specialized to elongate for signal transmission, neurons lose their regenerative capacity *(3)*. The morphological changes induced by intracellular tension relaxation potentially reverse this process, suggesting that the distinctive response to mechanical signals may play a crucial role in neurodevelopment. Mechanical force directly modulates the developmental state of neural cells, encouraging a neuro-specific cue for targeted neural injury therapies.

Meanwhile, we revealed the anti-inflammation and pro-regeneration in microglia responding to intracellular tension relaxation, although we did not explore the detailed mechanotransduction, which have been reported extensively, including YAP nuclear translocation and mechanosensitive ion channels *(14, 65)*. These findings about neuro- specialized immune cell provide wider insights within the central nervous system into the role of mechanical signals in regulating immune responses, and offer additional approaches for mitigating secondary inflammatory damage following SCI.

## Materials and Methods

### Synthesis and purification of enantiomers

Fmoc-^D^F^D^F^D^D and Fmoc-^L^F^L^F^L^D enantiomers were synthesized on Wang resin by standard 9-fluorenylmethoxycarbonyl (Fmoc) solid phase peptide synthesis. Blank Wang resin (2 g, Adamas) was activated in dimethylformamide (DMF, Sinoreagent) for 30 min, then treated with 5 equivalents each of D/L-aspartic acid (Asp, Adamas), 4- dimethylaminopyridine (DMAP, Adamas), and N, N’-diisopropylcarbodiimide (DIC, Adamas). After reacting for 3 hours at room temperature, the resin was washed five times with DMF. The Fmoc group was then deprotected using 20% v/v piperidine in DMF, which was examined using ninhydrin testing (Sinoreagent). Next, 3 equivalents of D/L-phenylalanine (Phe, Adamas), O-benzotriazole-N, N, N’, N’-tetramethyl-uronium- hexafluorophosphate (HBTU, Adamas), 1-hydroxybenzotriazole (HOBt, Adamas), and 10 equivalents of N, N-diisopropylethylamine (DIEA, Sigma-Aldrich) in DMF were added and reacted at room temperature for 40 min, followed by washing and Fmoc group deprotection. This cycle was repeated to synthesize the tripeptide without removing the Fmoc group of the final Phe. The peptide was cleaved from the resin using a standard solution of 95% trifluoroacetic acid (TFA, Adamas), 2.5% triisopropylsilane (TIS, Adamas), and 2.5% water. The products were purified using reversed phase high performance liquid chromatography (RP-HPLC, Agilent 1100, Fig. S1A) on an Inertsil ODS-SP-C18 column (GL Sciences). The purity of Fmoc-^D^F^D^F^D^D and Fmoc-^L^F^L^F^L^D detected using electrospray ionization mass spectrometry (ESI-MS, ZQ2000, Waters, Fig. S1B) was >95%, respectively. The ^1^H Nuclear Magnetic Resonance (NMR, Bruker Avance II-400 MHz) spectra results are as follows (10 mg/mL, DMSO-d6, δ, ppm, Fig. S1C): 2.58, 2.81 (dd, 2H, CH_2_); 2.69 (m, 2H, CH_2_); 2.92, 3.06 (dd, 2H,CH_2_); 3.94-4.22 (m, 4H, CH); 4.57 (m, 2H, CH_2_); 7.10-7.32 (m, 12H, Ar-H); 7.40 (td, 2H, Ar-H); 7.52 (d, 1H, NH); 7.60 (dd,2H, Ar-H); 7.87 (d, 2H, Ar-H); 8.11-8.39 (d, 2H, NH). The two enantiomers of the tripeptide were lyophilized and stored at -20 LJ.

### Self-assembled supramolecular hydrogel preparation

#### For material characterization and in vitro experiments

The lyophilized tripeptides were dispersed in phosphate buffer saline (PBS, pH 10), sonicated for 30 min to dissolve, and then mixed with an equal volume of PBS (pH 5). Hydrogel formed when pH was adjusted to 7.4 with 1 M NaOH, and then was employed to characterization. For cell culture, 200– 800 μL of hydrogel (1 mg/mL) was added to tissue culture-treated 24-well, 12-well, or 6- well polystyrene culture plates (Corning) or 35 mm glass-bottom dishes (Nest), air-dried at room temperature, and gently rinsed with ultrapure water to remove precipitated salt. Plates were sterilized under UV light for at least 30 min before cell culture.

#### For in vivo experiments

Lyophilized tripeptide powder was dissolved in sterile PBS (pH 7.4) at a concentration of 4 mg/mL. The solution was ultrasonically dispersed, heated to 90 °C for 30 min, and then rapidly cooled in an ice bath for the tripeptides to self-assemble into hydrogels (D-gel & L-gel).

### Molecular dynamics (MD) simulation

The two conformations of Fmoc-FFD enantiomers were constructed and optimized using Gaussian 09 software. Geometric optimization was performed with Density Functional Theory (DFT) employing B3LYP functional and 6-311+G(d,p) basis set, incorporating the polarizable continuum model to account for the influence of water. The force field simulation was performed using GROMACS 2022 software. The GAFF and Amber99sb- ildn force field were selected for Fmoc-FFD and proteins, respectively. The proteins or polymer system were solvated in cubic boxes of TIP3P water molecules situated at a minimal distance of 1.2 nm from the box edges. Sodium or chloride ions are added to the system to achieve neutralization. The cutoffs of the van der Waals interactions were set to be 1 nm, and the long-range electrostatic interactions were calculated by the particle mesh Ewald (PME) algorithm with a mesh spaced 0.14 nm. The neighbor list for the nonbonded interactions was updated every 10 steps. Hydrogens bonds were constrained using the LINCS algorithm and the time step for all simulations was 2 fs. The steepest descent method was employed to reduce the energy of 50,000 steps, followed by a 100 ps canonical ensemble (NVT) equilibration, a 100 ps constant-pressure and constant-temperature (NPT) equilibration. The temperature was maintained at 298.15 K with Velocity Rescale Thermostat, and the pressure was set at 1 bar with Parrinello-Rahman barostat throughout the duration of each simulation. Subsequently, a 100 ns MD production run was conducted, and the interaction energy (electrostatic interaction energy, E_Coul_, and Van der Waals interaction energy, E_VdW_) and hydrogen bonding (H-bond) were calculated to assess the stability of the system.

### Single-cell force spectroscopy (SCFS)

Astrocytes were seeded on gel-coated 6-well and cultured for 2 days. The adhesion force between single cell and the culture substrate was measured using FluidFM BOT system (Cytosurge AG) with a micropipette cantilever (4 μm aperture, nominal spring constant 0.3 N/m) featuring microfluidic channels. Before the measurements, the spring constant and deflection sensitivity of the cantilever were calibrated using the built-in Cytosurge software protocol. The system was mounted on an inverted microscope (iX 83, Olympus) to target individual cells for detachment. The cantilever approached cells at 10 μm/s with a preset force of 100 nN. Upon contact, a -500 mbar pressure was applied for 3 s to ensure tight cell-cantilever attachment, followed by cell detachment at a speed of 2 μm/s under - 500 mbar pressure, with a separation distance of 150 μm. Force-distance curves were recorded for each single cell.

### Förster resonance energy transfer (FRET)

Cells were seeded in gel-coated glass-bottom dishes and cultured for 2 days. The medium was replaced with serum-free DMEM at 2 h prior to transfection. Using Lipo3000 transfection reagent (GlpBio), the FRET-based actin probe plasmids (Actin-cpstFRET-Actin) were transfected into the cells at a concentration of 2.5 μg plasmid DNA per dish. After transfection for 5 h, the medium was replaced with complete culture medium, and cells were cultured for an additional 24 hours. FRET signal was observed using a laser scanning confocal microscope (LSM880, Zeiss). Single cell FRET ratio was analyzed using Zen Lite software (Zeiss). Actin-cpstFRET-Actin was a gift from Fred Sachs (Addgene plasmid #80643; http://n2t.net/addgene:80643; RRID:Addgene_80643).

### Cell morphological modeling

Astrocytes cultured on enantiomeric hydrogels for 2 days were processed with a 1,1’- dioctadecyl-3,3,3’,3’-tetramethylindocarbocyanine perchlorate (Dil, Beyotime) to stain cell membrane and 4’,6-diamidino-2-phenylindole (DAPI, Solarbio) to stain nucleus. After fixation, cells were imaged using a laser confocal microscope (SpinFV-COMB, Olympus) in Z-stack mode. The imaging files were imported into Imaris software for 3D image rendering, segmentation, and calculation of cellular and nuclear volumes.

### Fluorescence recovery after photobleaching (FRAP)

Astrocytes were stained with the Dil fluorescent probe (Beyotime) after cultured on hydrogels for 2 days. Imaging was conducted using a laser scanning confocal microscope (SpinFV-COMB, Olympus). A defined region of interest (ROI) was photobleached for 1 s using a 561 nm laser. Fluorescence recovery in the bleached area was monitored at low laser power (0.4%) for 148 s, with images acquired every 3 seconds. The recovery curves were normalized based on background fluorescence and overall photobleaching. Data were fitted using the cellSens Dimension software (Olympus) based on a classical exponential recovery model to calculate the fluorescence recovery halftime *τ* _1/2_.

### Fluorescence lifetime imaging microscopy (FLIM)

Astrocytes were seeded in gel-coated glass-bottom dishes and labeled with Flipper-TR^®^ probe (Cytoskeleton) following the manufacturer’s instructions. Flipper-TR^®^ is a fluorescent probe that specifically targets the plasma membrane and reports changes in membrane tension through its fluorescence lifetime variation, using FLIM (SP8 FALCON, Leica). Photon histograms from ROI were fitted with a double-exponential decay model to extract two decay times, τ_1_ and τ_2_. The higher fit amplitude decay time τ_1_ was used to characterize membrane tension, where a longer fluorescence lifetime indicates increased membrane tension.

### Animal surgery and injection procedure

All animal housing and experimental operations were approved by Experimental Animal Research Ethics Committee of Sichuan University (Approval No. KS20240250). In total, 75 rats were randomly assigned into three groups for subsequent injections: D-gel, L-gel, and Saline (pH 7.4 PBS); n=25 per group. All surgical procedures were performed by the same researchers. Female rats weighing 200–220 g were anesthetized with 2.5% isoflurane in oxygen-enriched air and placed in a prone position on a dissecting board. The paraspinal muscles were bluntly dissected to expose the T9-T11 vertebrae. A laminectomy was performed to expose the spinal cord. Severe spinal cord compression injury was induced by transversely compressing the T10 spinal cord completely for 10 seconds using forceps (Dumont, 028-2SP-PO). Subsequently, Treatment agents were administered at SCI site using a using a sterile 30G insulin syringe (OMNICAN) with a total volume of 15LJμL injection per rat. After injection, the needle was slowly withdrawn, ensuring the dura mater remained intact with no fluid leakage. The muscles and skin were then sutured separately, and a penicillin ointment was applied to the sutured area to prevent bacterial infection. Manual bladder emptying was performed twice daily for 7 days. Rats were administered meloxicam (dissolved in 5% glucose solution, 1 mg/kg, QCS) via oral gavage once daily for 3 days.

### Single-nucleus RNA sequencing (snRNA-seq)

For snRNA-seq, fresh tissue samples were collected on postoperative day 10 and immediately placed in cold PBS to prevent RNA degradation. Tissue was mechanically dissociated into small pieces and then incubated with a dissociation buffer containing papain to release the nuclei. Nuclei were isolated through filtration using a cell strainer, followed by centrifugation to pellet the nuclei. The nuclei were resuspended in an appropriate buffer, and their concentration was quantified using a hemocytometer or automated cell counter. After confirming the integrity of the nuclei, single-nucleus suspensions were loaded onto a droplet-based platform (10x Genomics Controller) for high-throughput sequencing. RNA was captured and reverse transcribed to cDNA, which was then amplified using PCR. The resulting libraries were subjected to quality control and assessed for size distribution and purity. Sequencing was performed on a high-throughput platform (NovaSeq 6000, Illumina) to generate transcriptomic data at the single-nucleus level. Data analysis involved the alignment of sequencing reads to the reference genome, followed by quality filtering, normalization, and clustering of individual nuclei based on gene expression profiles. Differential gene expression analysis was performed to identify genes with significant expression changes across different conditions. Pathway enrichment analysis was conducted to identify biological processes and pathways associated with the observed expression patterns.

### Statistical analyses

Data was analyzed using GraphPad Prism 10.1 software and presented as means ± standard deviation (SD) unless otherwise noted. The comparisons between two groups were performed using an unpaired Student’s two-tailed t-test. where the t-value was computed as the difference in means divided by the standard error of the difference. The degrees of freedom for the unpaired t-test were calculated as *df=n*_*1*_ *+ n*_*2*_ *– 2*, where n_1_ and n_2_ are the sample sizes of the two groups being compared. Differences between three or more groups were analyzed using one-way analysis of variance (ANOVA), followed by Tukey’s post-hoc test for multiple comparisons. The F-value was calculated by dividing the variance between groups by the variance within groups. The corresponding degrees of freedom were determined as *df*_*between*_*=k* − *1* and *df*_*within*_*=N*−*k*, where *k* represents the number of groups and *N* the total sample size. A p-value < 0.05 was considered statistically significant.

## Acknowledgments

The authors thank Dr. Siying Qin in the Core Facilities (School of Life Sciences, Peking University), Xi Hu and Shiqiang Wang (Quantum Design China), Yin Liu (Department of science and technology development, Southwest University), Junshu Zhu and Lin Wu (Integrative Science Center of Germplasm Creation in Western China (CHONGQING) Science City), for their assistance with FluidFM BOT. We thank Wenwu Wang (College of Materials Science & Engineering, Sichuan University) for his help of AFM capturing and analysis. The authors would like to thank Jiao Lu and Guolong Meng (College of Biomedical Engineering, Sichuan University), for their assistance with laser confocal microscopy. We are grateful to Dr. Cheng Qiao (ZEISS), Mrs. Jing Tang (Olympus), Junyang Shi (Olympus), Liyuan Wu (Olympus), for their technical support with confocal imaging. We would like to thank Linzhu Li in the Life Science Core Facilities (College of Life Sciences, Sichuan University) for help of digital scanning system and laser confocal microscopy. We thank Prof. Xianwei Meng and Dr. Shimei Li (Technical Institute of Physics and Chemistry, Chinese Academy of Sciences) for help of cell culture. We also thank Prof. Jiang Wu and Dr. Jirui Wen for their support on flow cytometry. We thank Personal Biotechnology Co., Ltd. (Shanghai, China) for their support in data analysis for the single-cell sequencing. We thank ChatGPT (OpenAI) for providing language editing support and improving the grammatical accuracy of this paper.

## Author contributions

† These authors contributed equally to this work. Conceptualization: YL, QA; Methodology: YW, YL, XF, HZ, RL, XL, XL; Supervision: QA, ZH, GY, AW Visualization: YW, YL, RC, ZY; Writing—original draft: YL, YW; Writing—review & editing: YL, QA, YW.

## Data and materials availability

All data are available in the main text or the supplementary materials.

## Supplementary Materials

This file includes:

Materials and Methods

Supplementary Text

Fig. S1 to S28

