## Supplementary materials for "Enantiomeric Hydrogel-Manipulated Mechanotransduction Triggers Neurogenesis and Immunomodulation for Spinal Cord Repair"

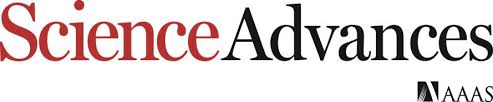

Supplementary Materials for

**Enantiomeric Hydrogel-Manipulated Mechanotransduction Triggers Neurogenesis and Immunomodulation for Spinal Cord Repair**

Ya Li†, Yulin Wang† *et al.*

**This PDF file includes:**

Materials and Methods

Supplementary Text

Figs. S1 to S28

**Materials and Methods**

**1. Characterization of tripeptide enantiomers and supramolecular hydrogels**

*Implementation and analyses of UV-vis spectroscopy and Circular dichroism (CD) spectroscopy*

The UV-vis spectra of enantiomers (Fmoc-^D^F^D^F^D^D and Fmoc-^L^F^L^F^L^D, 1 mg/mL in 50%:50% methanol:acetonitrile solution) and their corresponding self-assembled hydrogels (D-gel and L-gel, 1 mg/mL in PBS) were recorded using a spectrophotometer (UV3600, Shimadzu) with a 1 cm path-length quartz cuvette (Starna) at continuous scan mode (step: 1 nm). As shown in Fig. 1B, the characteristic peak at 215 nm corresponded to the π → π* transition of the conjugated system in the amide region (C=O and N-H). Peak at 255–267 nm corresponded to the π-π* transition of the phenyl group in phenylalanine, while peaks at 290 nm and 301 nm are attributed to the conjugated system of the Fmoc group.

The CD spectra of the same samples were next recorded using a spectropolarimeter (Chirascan Plus V100, Applied Photophysics) and the signal intensity was calibrated based on UV-vis absorbance value. As shown in Fig. 1B, the peptide enantiomers exhibited three peaks with one at 215 nm corresponding to the amide bond and other two at 228 nm and 237 nm attributed to phenyl group. The hydrogels exhibited a characteristic Cotton effect, with speaks at 195 nm and 218 nm indicating the formation of β-sheet structures and peaks at 228 nm, 237 nm, and 254 nm assigned to π → π* transitions and π-π stacking of phenyl rings. The broad peaks between 275–305 nm contributed to extended conjugation between the Fmoc group and phenyl rings. These results reflect the highly ordered stacking of internal structures and interlaced aromatic ring systems within the hydrogels.

*Transmission electron microscopy (TEM)*

The dilute gel solution (0.1 mg/mL in PBS) was dropped on a glow-discharged grid (300-mesh copper grid with carbon support film, Ted Pella), and the excess liquid was absorbed using filter paper. The gel was stained with 2 wt% uranyl acetate and air-dried subsequently. The negatively stained samples were observed using a TEM (JEM-1400, JEOL) at 120 kV.

*Atomic force microscopy (AFM)*

The dilute gel solution (0.2 mg/mL in PBS) was dropped onto a freshly cleaved mica disk (Ted Pella), air-dried at room temperature, and gently rinsed three times with ultrapure water. Imaging was performed using an AFM (Multimode 8, Bruker).

*Implementation and analyses of Fourier transform infrared spectroscopy (FT-IR)*

The hydrogels (2 mg/mL in PBS) were dropped onto a CaF_2_ window, dried under an infrared lamp. The FT-IR spectra were recorded on a spectrometer (Nicolet iS50, Thermo Fisher). The spectra are identical between D-gel and L-gel (Fig. S2A), validating the consistency of their chemical groups. The observed peaks are attributed to the following: 3297 cm^-1^ (O-H stretching of H_2_O and N-H stretching of amide I region), 3064 cm^-1^ (=C-H stretching of aromatic rings), 1713 cm^-1^ (C=O stretching of -COOH), 1690 cm^-1^ and 1646 cm^-1^ (C=O stretching of amide I region), 1535 cm^-1^ (N-H bending and C-N stretching of amide II region), 1450 cm^-1^ (C=C skeletal stretching of aromatic rings), and 1256 cm^-1^ and 1222 cm^-1^ (coupled C-N stretching and N-H bending vibrations of amide III region).

*X-ray scattering*

The hydrogels (4 mg/mL) were analyzed using synchrotron small-angle X-ray scattering (SAXS) and wide-angle X-ray scattering (WAXS) detectors (PILATUS4, DECTRIS), with a scattering vector range of 0.011 < q < 2.6 Å^-1^ and PBS as background. The 2D scattering data was processed using Igor Pro software to generate 1D intensity vs. wave vector plots.

*Rheological measurements*

Rheological properties of the hydrogels were characterized using a rheometer (MCR302, Anton Paar) at 37 °C. The viscoelastic behavior was evaluated under frequency sweep mode over 0.1–10 Hz. The stress relaxation behavior was examined in the frequency range of 0.1–10 Hz under 10% strain, and the resulting stress relaxation data were modeled and normalized using the viscoelastic Maxwell-Weichert model. The viscoelasticity variation responding to cerebrospinal fluid (Solarbio) was examined under time sweep mode at 1 Hz.

**2. Cell culture**

*Astrocyte culture*

Mouse astrocyte cell line C8-D1A was cultured in Dulbecco’s modified Eagle medium (DMEM, Gibco) supplemented with 10% fetal bovine serum (FBS, Gibco) and 1% penicillin-streptomycin (Gibco) at 37 °C in a humidified incubator with 5% CO_2_ (Thermo Fisher). The medium was replaced every two days, and cells were passaged with EDTA-free trypsin (Gibco) at 70–80% confluency.

*Microglial cell culture*

Mouse microglial cell line BV2 was cultured under the same conditions as astrocytes. BV2 cells exhibit semi-suspension growth and were collected through gently pipetting.

**3. *In vitro* experiments**

*Cell viability*

Astrocytes and microglia cells (1×10^5^ cell/mL) were seeded on different substrates (D-gel, L-gel, and standard adhesion culture-treated substrate, 24-well plates) and cultured for 1, 3, and 5 days. For CCK-8 assays, 10% (v/v) CCK-8 solution (APE×BIO) was added to each well and incubated at 37 °C in the dark for 2 h. The absorbance of formazan at 450 nm was measured using a microplate reader (synergy H1, BioTek), and viability was calculated as the percentage of that of the control group. For live/dead staining assays, cells were washed with PBS and stained using the Calcein-AM/Propidium Iodide (PI) live/dead cell staining kit (Solarbio) according to the manufacturer’s instructions. Stained cells were imaged under an inverted fluorescence microscope (Ti2-U, Nikon).

*Cell proliferation*

Cells cultured on D-gel and L-gel for 2 d were labeled using EdU Imaging Kit (APE×BIO) and EdU Flow Cytometry Assay Kit (APE×BIO) according to the manufacturer's protocols. Labeled cells were imaged under an inverted fluorescence microscope (Ti2-U, Nikon), and the proliferation rate was quantitatively analyzed using a flow cytometer (NovoCyte, Agilent).

*Colony formation*

Astrocytes (500 cells/well) were seeded on D-gel and L-gel (6-well plates) and cultured for 7 d. Cells were fixed with 4% paraformaldehyde (PFA, Biosharp) for 20 min and stained with crystal violet solution (Solarbio) for 30 min, then observed under a microscope (APX100, Olympus).

*Apoptosis*

Cells cultured on chiral gel-coated 6-well plates for 2 d were stained using Annexin V-FITC/PI apoptosis kit (APExBIO) according to the manufacturer’s protocol. Labeled cells were analyzed with a flow cytometer (NovoCyte, Agilent). Cells treated with hydrogen peroxide (200 μM, overnight) were used as positive control of Annexin V-FITC, and cells treated with 20% (v/v) hydrogen peroxide for 5 minutes served as positive control of PI.

*Mitochondrial membrane potential (MMP)*

The MMP variation was detected using JC-10 fluorescent probe (Biosharp). At high MMP, JC-10 aggregates in the mitochondrial matrix, emitting red fluorescence, while at low MMP, it exists as monomers outside the mitochondria, emitting green fluorescence. Cells were seeded on gel-coated glass-bottom dishes and stained with JC-10 MMP kit according to the manufacturer’s instructions. Live-cell imaging was performed using a laser confocal microscope (LSM880, Zeiss).

*Intracellular ATP level*

Cells were seeded in gel-coated 12-well plates and cultured for 48 h, and then were collected and lysed through ultrasonic for 1 min on ice. Intracellular ATP level was determined using a micro-assay method based on creatine kinase-catalyzed phosphocreatine produced from ATP. The phosphocreatine content was quantified using phosphomolybdic acid colorimetric method using a microplate reader (synergy H1, BioTek). Total protein concentration was determined using a BCA protein assay kit (Beyotime) for normalization. ATP concentration was calculated using the formula: CATP (μmol/mg protein) = 2 × ΔA_samples_ / (ΔA_standard_ × C_protein_).

*Cell migration*

Astrocytes were seeded on gel-coated 6-well with Ibidi Culture-Inserts and cultured until reaching 100% confluency. The inserts were then removed to create a neat cell-free area on the gel surface. The cell sprawling into this area were monitored over 24 h using a live-cell imaging system (APX100, Olympus), and the migration rate was quantified.

*RNA-sequencing*

Astrocytes were seeded in gel-coated 12-well plates until reaching 80% confluence. Total RNA was extracted using Trizol reagent (Invitrogen). RNA concentration, quality, and integrity were assessed using a NanoDrop 2000 spectrophotometer (Thermo Fisher) and 2100 Bioanalyzer system (Agilent). The cDNA library was constructed by PersonalBio and sequenced on the NovaSeq 6000 platform (Illumina). RNA sequencing data were trimmed using cutadapt to remove poly(A) tails and low-quality reads, and subsequently aligned to the Mus musculus GRCm38 genome (Ensembl). Gene-level counts were summarized using HTSeq, and gene expression values were calculated as fragments per kilo bases per million fragments (FPKM) with the addition of log2 transformation and pseudo-count of 1. Heatmaps of gene expression were generated using the pheatmap R package. Pairwise differential gene expression analysis was conducted using the DESeq R package, with a threshold of fold change > 2 and p-value < 0.05. Volcano plots of differentially expressed genes were created using the ggplot2 R package. GO and KEGG enrichment analyses were performed using topGO and KAAS, respectively. The filtering criteria were set to fold change > 2 and p-value < 0.05. Results were visualized as GO enrichment bar plots and KEGG pathway diagrams.

*Blebbistatin processing*

Astrocytes were seeded in standard tissue culture treated 96-well plates until reaching 70% confluence. The medium was then replaced with blebbistatin contained (0.1-16 μM) complete DMEM and the cells were cultured for 24 h. The cell viability was measured through CCK-8 assay.

Astrocytes were cultured in standard tissue culture treated 12-well plates until reaching 80% confluence. The cells were treated with 4 μM blebbistatin for 24 h, and then labeled using EdU Flow Cytometry Assay Kit (APE×BIO). The proliferation rate was quantitatively analyzed using a flow cytometer (NovoCyte, Agilent).

*Calcium ion staining*

Astrocytes were cultured in 24-well plates coated with gels. Intracellular calcium ions were stained using a Fluo-4 calcium assay kit (Beyotime) according to the manufacturer’s instructions. Fluo-4 bound to intracellular calcium ions and emitted green fluorescence upon excitation, which is observed using an inverted fluorescence microscope (Ti2-U, Nikon).

**4. *In vivo* experiments**

*Animal inclusion and exclusion criteria*

At 24 h after surgery, all rats were evaluated for hind limb motor function in an open field. Rats exhibiting any hind limb movement were excluded from the study and euthanized by exposure to an overdose of carbon dioxide.

*Anterograde BDA spinal tracing*

At 14 days after SCI, rats were anesthetized and secured on a stereotaxic apparatus. A laminectomy was performed 2 cm rostral to the injury site to expose the spinal cord. Two injection coordinates were identified on the two side of the central spinal blood vessel (±0.5 mm, ±1 mm), with two injection depths for each coordinate (-0.8 mm and -1.2 mm). A 5 μL glass micro-syringe was used to inject 0.5 μL of 10% biotinylated dextran amine (BDA, Mw=10,000 Da, Invitrogen) at each coordinate. Following the injections, the rats were maintained until day 28 post-SCI. Tissue sections were processed with Cy3-conjugated streptavidin (APE×BIO) to visualize BDA-labeled axons.

*Functional recovery evaluation*

The motor function recovery of hindlimb in rats was assessed using the Basso, Beattie, and Bresnahan (BBB) scoring scale, which ranges from 0 (no motor function) to 21 (normal motor function). Rats were placed in the open field, and two trained experimenters independently scored the locomotor behavior of rats. The locomotor activity of rats was further evaluated with open field test. Rats were placed individually in a 100 cm × 100 cm arena for 5 min. Locomotion behavior was assessed using a video tracking system (OFT-200, TECHMAN), recording parameters including locomotion velocity and activity time. The arena was cleaned with 70% ethanol between tests to avoid olfactory interference. An intelligent gait analysis system (GAT-100, TECHMAN) was used to record rat locomotion, with a 100 cm long, adjustable-width runway and a high-resolution camera. Rats were placed on the runway and recorded at postoperative day 7, 14, 21, and 28. Sensory function recovery was assessed using the tail-flick pain test. After securing the rats, an infrared heat radiator (PL-200, TECHMAN) was applied to the tails (2 cm from the base), and the latencies were recorded.

*Electrophysiological analyses*

Motor evoked potentials (MEPs) were assessed on postoperative day 28. After anesthesia, rats were placed in a prone position and stabilized. The scalp was incised to expose the skull, and bilateral 2-mm craniotomies performed at coordinates 1 mm anterior to bregma and 2 mm lateral to the sagittal suture using a bone drill (78001, RWD). A bipolar electrode was used to stimulate the motor cortex with single-pulse stimulation of 10 V intensity and 0.2 ms pulse width. A recording electrode was placed in the contralateral gastrocnemius muscle, and electrophysiological signals were recorded using a biological signal acquisition system. (RM6240E, Chengyi). The stimulation and recording points were maintained at consistent distances for all rats.

*Tissue collection and processing*

After anesthesia with sodium pentobarbital, blood cells were removed by cardiac perfusion with 0.9% saline, followed by perfusion with 4% paraformaldehyde (PFA, Biosharp) to fix the tissue. The spinal cords were then collected and further fixed overnight at 4 °C in 4% PFA. Then, the tissues were dehydrated through graded ethanol, cleared in xylene, and infiltrated with paraffin at 60 °C for 3–4 h. The tissues were then embedded in paraffin blocks, trimmed, and stored until sectioning. For frozen sections, the fixed spinal cords were immersed in 30% sucrose for 24 h, embedded in optimal cutting temperature (OCT) compound, and sectioned at 10 μm using a cryostat (Leica CM1950) for staining and analysis.

*High-throughput protein microarray technology*

The immune cytokine levels at the injury site on postoperative day 2 and 7 were analyzed using Luminex liquid suspension chip. Spinal cord tissues were collected after perfusion with 0.9% saline to remove blood cells and total protein was extracted using a BCA protein assay kit (Beyotime). The Bio-Plex Pro Rat Panel 23-plex Cytokine kit (Bio-Rad) was used following the manufacturer’s instructions. Briefly, protein samples were incubated in plate wells embedded with microbeads for 1 h, followed by a 30 min incubation with corresponding antibodies. Streptavidin-PE was then added to each well and incubated for 10 min. The signals were detected using the Bio-Plex 200 system (Bio-Rad).

**5. Biological assays for *in vitro* and *in vivo* experiments**

*Hematoxylin and eosin (H&E) staining*

Tissue sections were deparaffinized in xylene and rehydrated through graded ethanol to distilled water. Sections were stained with hematoxylin for 3–5 min, rinsed in water, differentiated briefly in 1% acid-alcohol, and blued in an alkaline solution. After rinsing, sections were counterstained with eosin for 1–3 min, dehydrated through graded ethanol, cleared in xylene, and mounted for microscopic observation.

*Immunofluorescence (IF) staining*

For both cell samples and tissue sections, samples were washed with PBS and fixed with 4% PFA. Permeabilization was performed using 0.2% Triton X-100 for 10–15 min, followed by blocking with 5% w/v bovine serum albumin (BSA, MCE) solution for 1 h. Samples were incubated with primary antibodies diluted in 3% w/v BSA at 4 °C overnight. After three PBS washes, secondary antibodies (1:200, SAB) were added and incubated at room temperature in the dark for 1 h. Nuclei were counterstained with DAPI (Solarbio). Additionally, cytoskeleton was stained with rhodamine-conjugated phalloidin (Sigma). Imaging was performed using a laser scanning confocal microscope (LSM880, Zeiss) for cell samples and a slide scanner (VS200, Olympus) for tissue sections. Fluorescence intensity quantification was performed using ZEN lite and Image J software.

*Western blot (WB)*

Cells and tissue samples were lysed using RIPA buffer (containing protease and phosphatase inhibitors) to extract total protein. The protein concentration was determined using BCA protein assay kit. Equal amounts of protein (20-40 μg) were separated by SDS-PAGE (Epizyme) and transferred to a PVDF membrane (Epizyme). The membrane was blocked with 5% w/v BSA in TBST (Epizyme) for 1 h at room temperature. After blocking, the membrane was incubated overnight at 4 °C with primary antibodies (diluted in TBST with 5% w/v BSA). Then, the membrane was washed three times with TBST and incubated with HRP-conjugated secondary antibodies (1:5000, SAB) for 1 h at room temperature. The membrane was then washed again with TBST and developed using an enhanced chemiluminescence (ECL, Epizyme) detection system. The protein bands were visualized and quantified using a chemiluminescence imager (ChemiDoc XRS+, Bio-Rad).

*Flow cytometry (FC)*

Cells were harvested and washed twice with PBS, then resuspended in 1% w/v BSA. Cells were incubated with specific dyes or fluorophore-conjugated antibodies for 30 min at 4 °C in the dark. After washing twice to remove unbound dyes or antibodies, the cells were resuspended in staining buffer for analysis. For intracellular protein staining, cells were fixed with 4% paraformaldehyde for 20 minutes, permeabilized with Saponin (Beyotime), and then incubated with fluorophore-conjugated antibodies. Stained samples were analyzed using a flow cytometer (NovoCyte, Agilent), and data were processed with NovoExpress software.

*Antibodies for IF, WB, and FC*

The following primary antibodies were used for in vivo and in vitro experiments. Rabbit anti-vinculin (1:200, SAB), rabbit anti-ITGB1 (1:200, SAB), rabbit anti-paxillin (1:200, SAB), rabbit anti-talin (1:200, SAB), rabbit anti-pMYH9 (1:200, SAB), rabbit anti-lamin A/C (1:200 for IF, 1:1000 for WB, SAB), rabbit anti-HDAC1 (1:200 for IF, 1:1000 for WB, SAB), rabbit anti-YAP (1:200 for IF, 1:1000 for WB, SAB), rabbit anti-piezo1 (1:200 for IF, 1:1000 for WB, SAB), rabbit anti-TRPV4 (1:100 for IF, 1:1000 for WB, Bioss), rabbit anti-TUBB3 (1:150, Proteintech), rabbit anti-GFAP (1:200, SAB), rabbit anti-MBP (1:200, SAB), rabbit anti-laminin b1 (1:150 for IF, Proteintech), rabbit anti-CHAT (1:150, Proteintech), rabbit anti-HTR2B (1:200, SAB), mouse anti-iba1 (1:500, GeneTex), rabbit anti-ARG1 (1:200, SAB), rabbit anti-iNOS (1:200, SAB), APC conjugated anti-MRC1 (1:200, Biolegend), FITC conjugated anti-iNOS (1:100, SAB), rabbit anti-β actin (1:5000 for WB, SAB), rabbit anti-laminin b1 (1:1000 for WB, Cell Signaling Technology).

**Supplementary Text**

In snRNA-seq, we identified 8 astrocyte clusters using unbiased clustering (Fig. 4A) and analyzed their gene characteristics through KEGG pathway enrichment. In cluster 1 with biggest population, endocytosis and autophagy pathways were upregulated and corresponding marker genes were highly specific (Fig. S18A), confirming to the characteristics of phagocytic reactive astrocytes. In cluster 2, besides endocytosis and phagosome, immune-related pathways emerged with corresponding marker genes (Fig. S18B), aligning with the characteristics of immune reactive astrocytes. In cluster 3, proliferation-related pathways were upregulated massively, suggesting that they are scar-forming astrocytes (Fig. S18C).

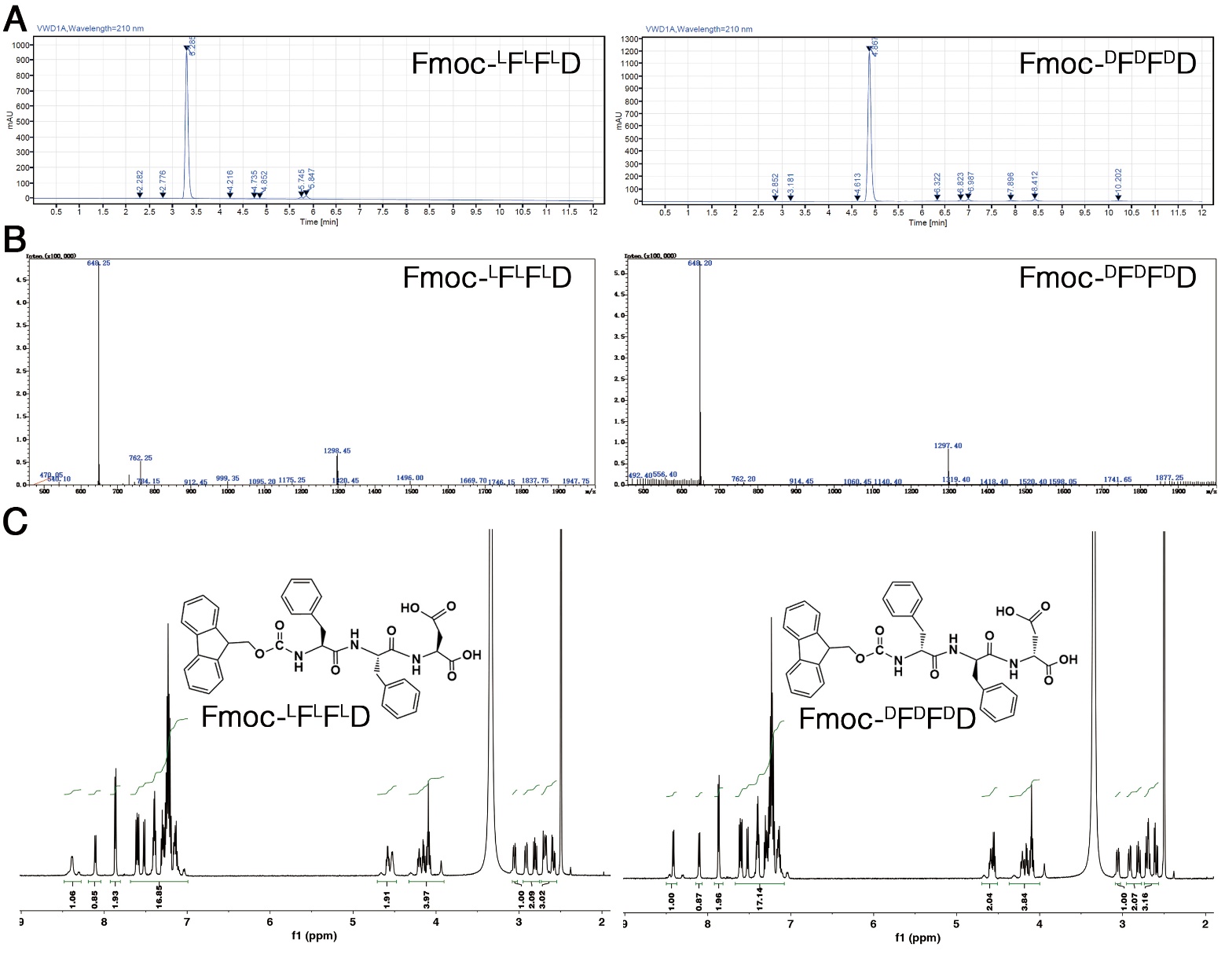

Fig. S1. Synthesis and characterization of Fmoc-FFD enantiomers. (A) HPLC profiles. (B) ESI-mass spectra. (C) ^1^H NMR spectra.

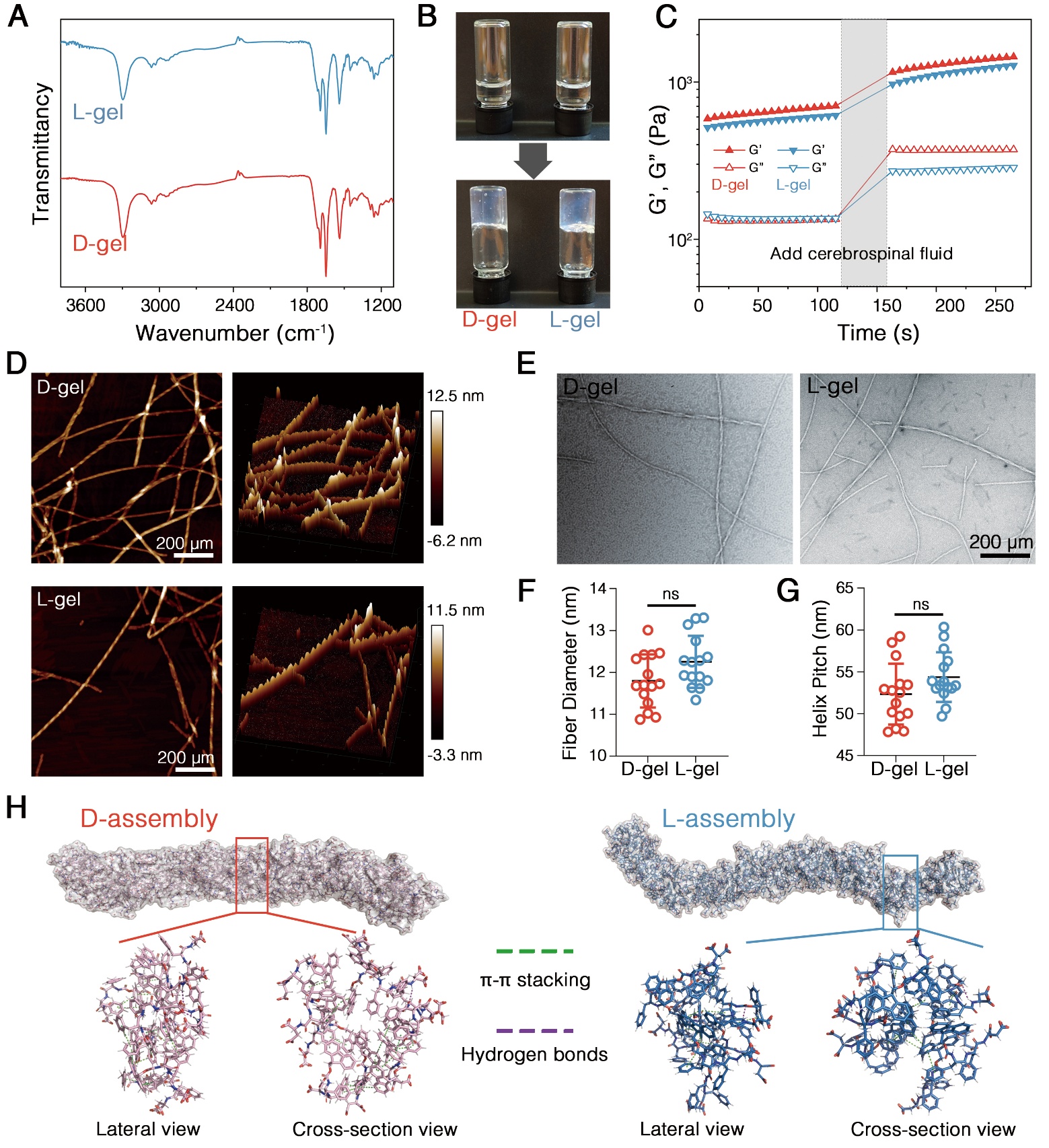

Fig. S2. Characterization of enantiomeric hydrogels. (A) FT-IR spectra. (B) Photos of enantiomeric tripeptides dissolved in DMSO (top) and self-assembled hydrogels in PBS (bottom). (C) The change of rheological strength of hydrogels responding to cerebrospinal fluid addition. (D) Representative AFM images of enantiomeric hydrogels in 2D (left) and 3D (right). (E) TEM images of enantiomeric hydrogels. (F, G) Quantification of fiber diameters and pitches of self-assembled structures based on TEM images. (H) Structural visualization of enantiomeric assemblies based on MD stimulation, showing filaments formation through π-π stacking (primary) and hydrogen bonding (secondary) of Fmoc-FFD molecules.

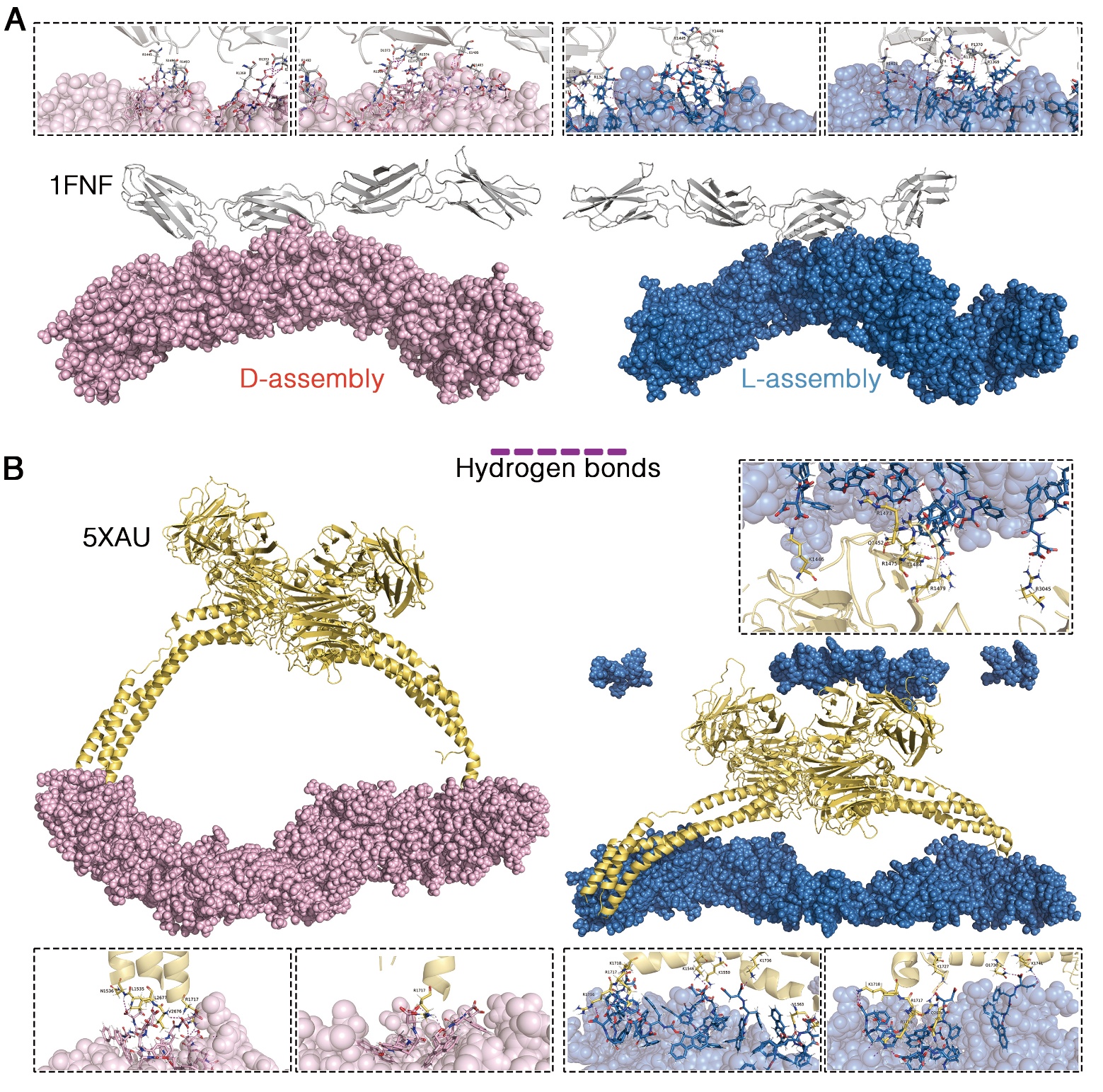

Fig. S3. Interactions between enantiomeric assemblies and two ECM-related proteins. (A) Interaction of the two assemblies 1FNF (gray), including a magnified view of the binding sites. 1FNF is a fragment of human fibronectin encompassing type-III repeats 7 through 10. (B) Interaction of the two assemblies with 5XAU (yellow), including a magnified view of the binding sites. 5XAU is the crystal structure of the integrin-binding fragment of laminin-511. Hydrogen bonding predominates in the interactions between the assemblies and proteins.

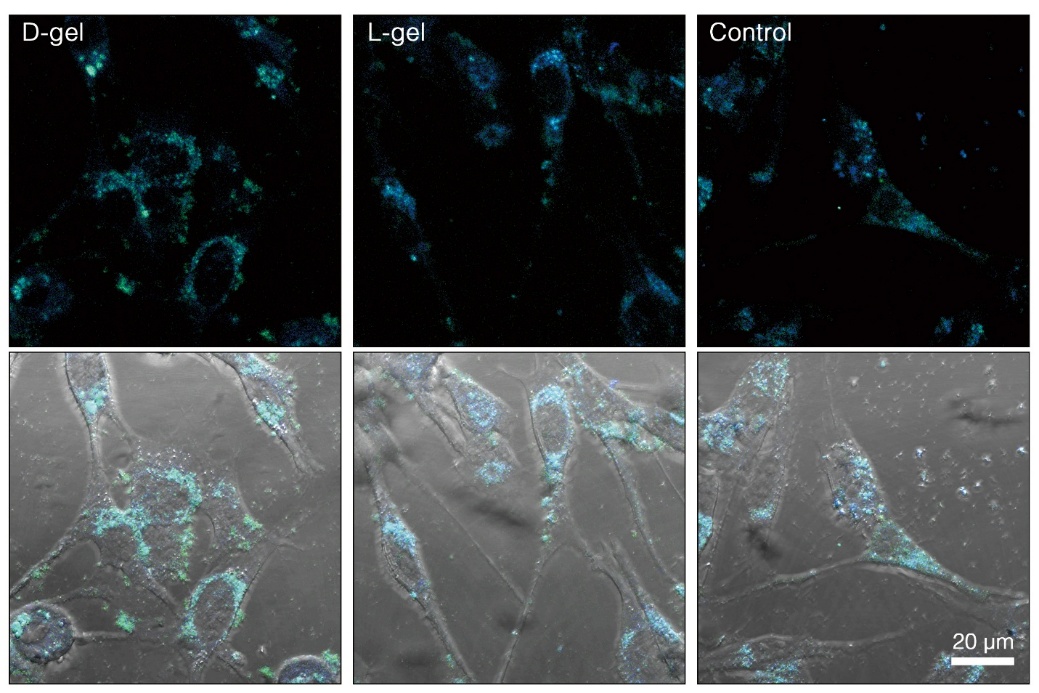

**Fig. S4.** Fluorescent micrographics of astrocytes transfected with Actin-cpstFRET-Actin plasmid. Changes in fluorescence color indicates intracellular tension variation.

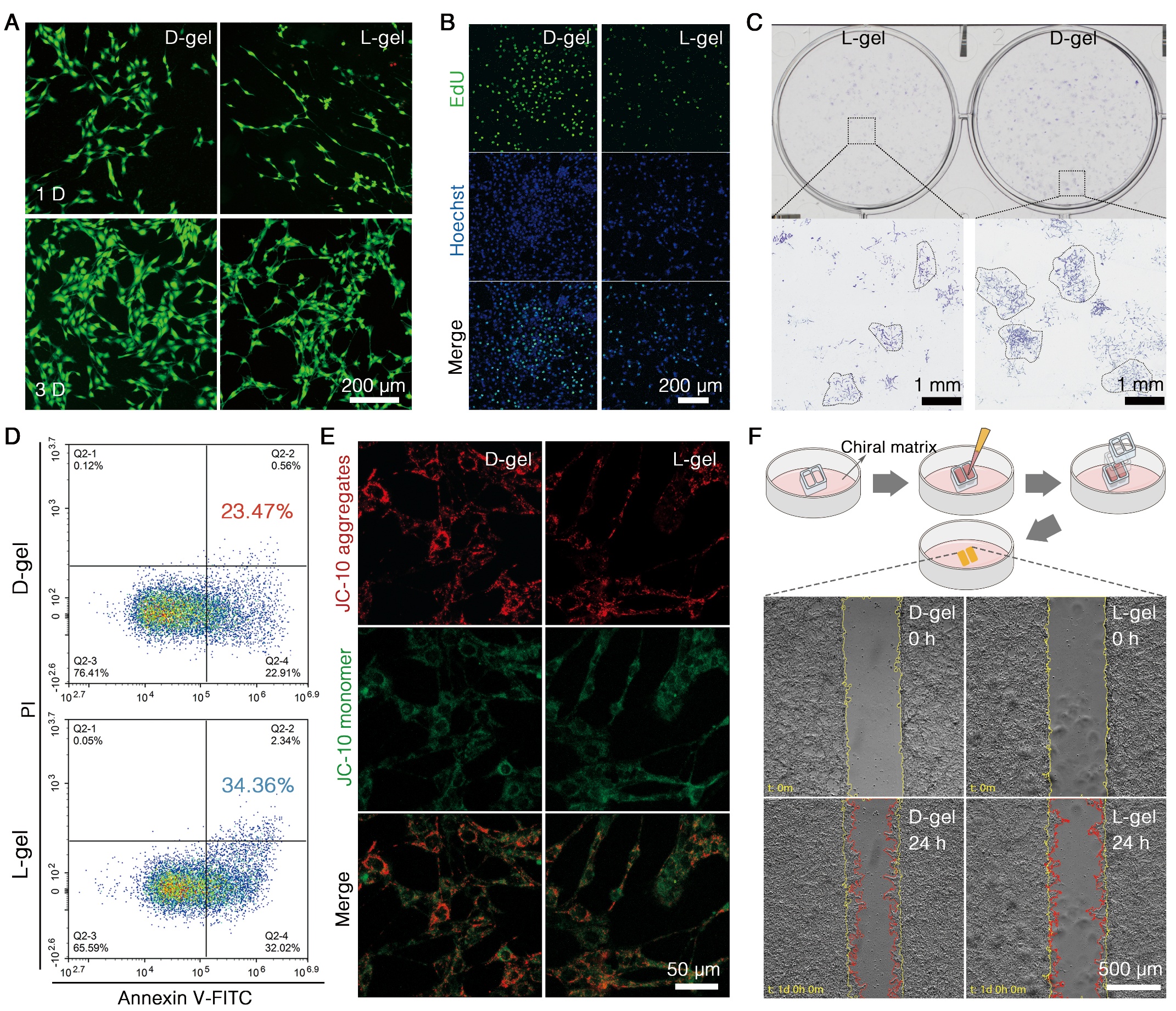

**Fig. S5. Effects of enantiomeric hydrogels on astrocyte behavior**s. (A) Representative live/dead staining fluorescent images of astrocytes cultured on enantiomeric hydrogels. (B) Representative fluorescent micrographics of EdU-stained proliferating astrocytes. (C) Representative images of crystal violet-stained astrocytes, exhibiting colony formation. (D) Representative percentage of apoptotic astrocytes (indicated by numbers, representing the sum of the second and fourth quadrants) determined by flow cytometric analysis with Annexin V–FITC/PI staining. (E) Representative fluorescent micrographics of JC-10-stained astrocytes, indicating mitochondrial membrane potential. (F) Migration of astrocytes on enantiomeric hydrogels. Yellow and red lines indicate cell edges before and after migration, respectively.

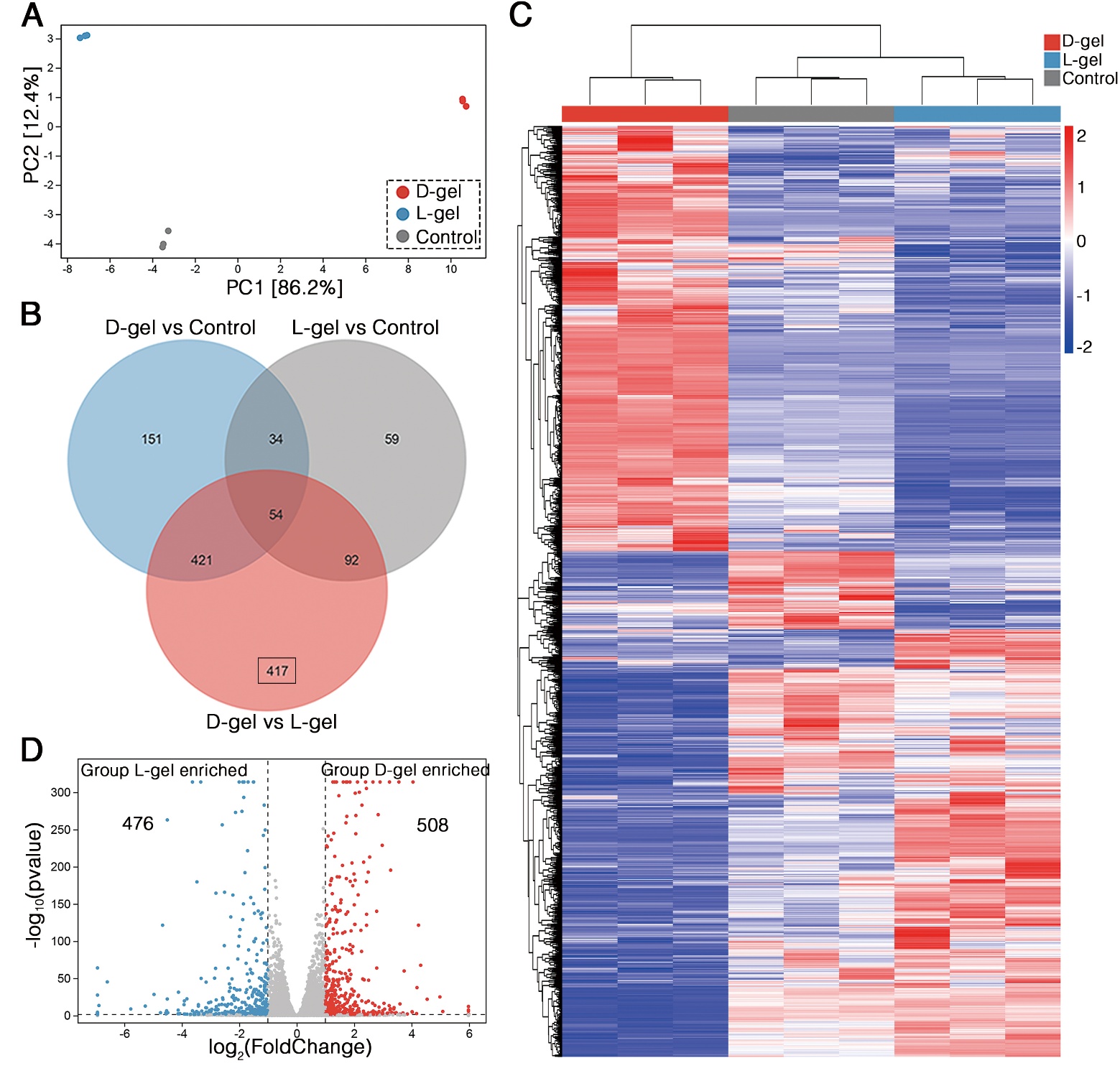

**Fig. S6. Transcriptomic analysis of astrocytes.** (A) Principal component analysis (PCA) of three groups. (B) Venn diagram of differentially expressed genes (DEGs) in pairwise comparisons between groups. (C) Volcano plot of DEGs between the D-gel and L-gel groups. (D) Clustering analysis of all DEGs across the three groups.

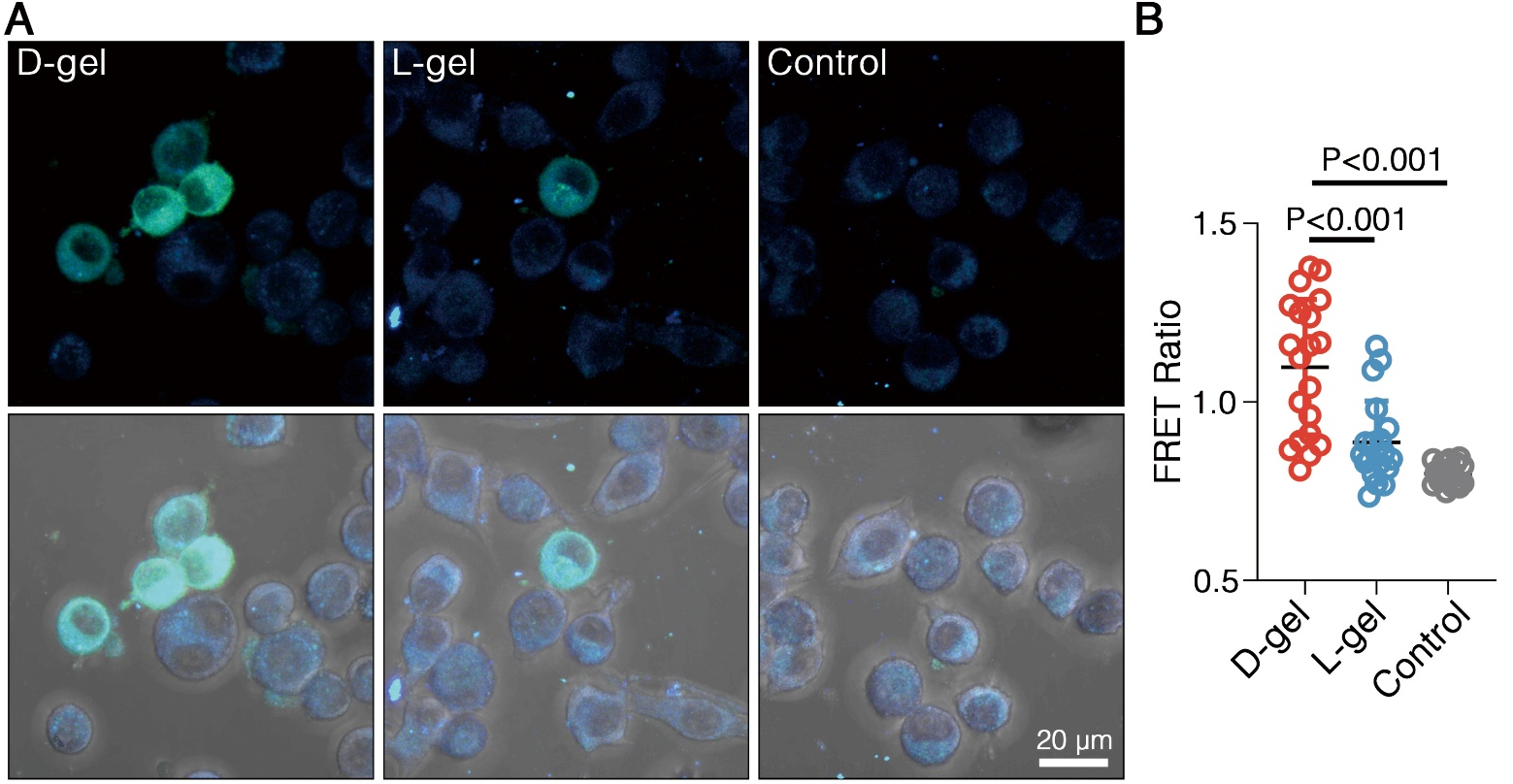

**Fig. S7.** (A) Fluorescent micrographics of microglia transfected with Actin-cpstFRET-Actin plasmid. Changes in fluorescence color indicates intracellular tension variation. (B) FRET efficiency of microglia cultured on enantiomeric hydrogels.

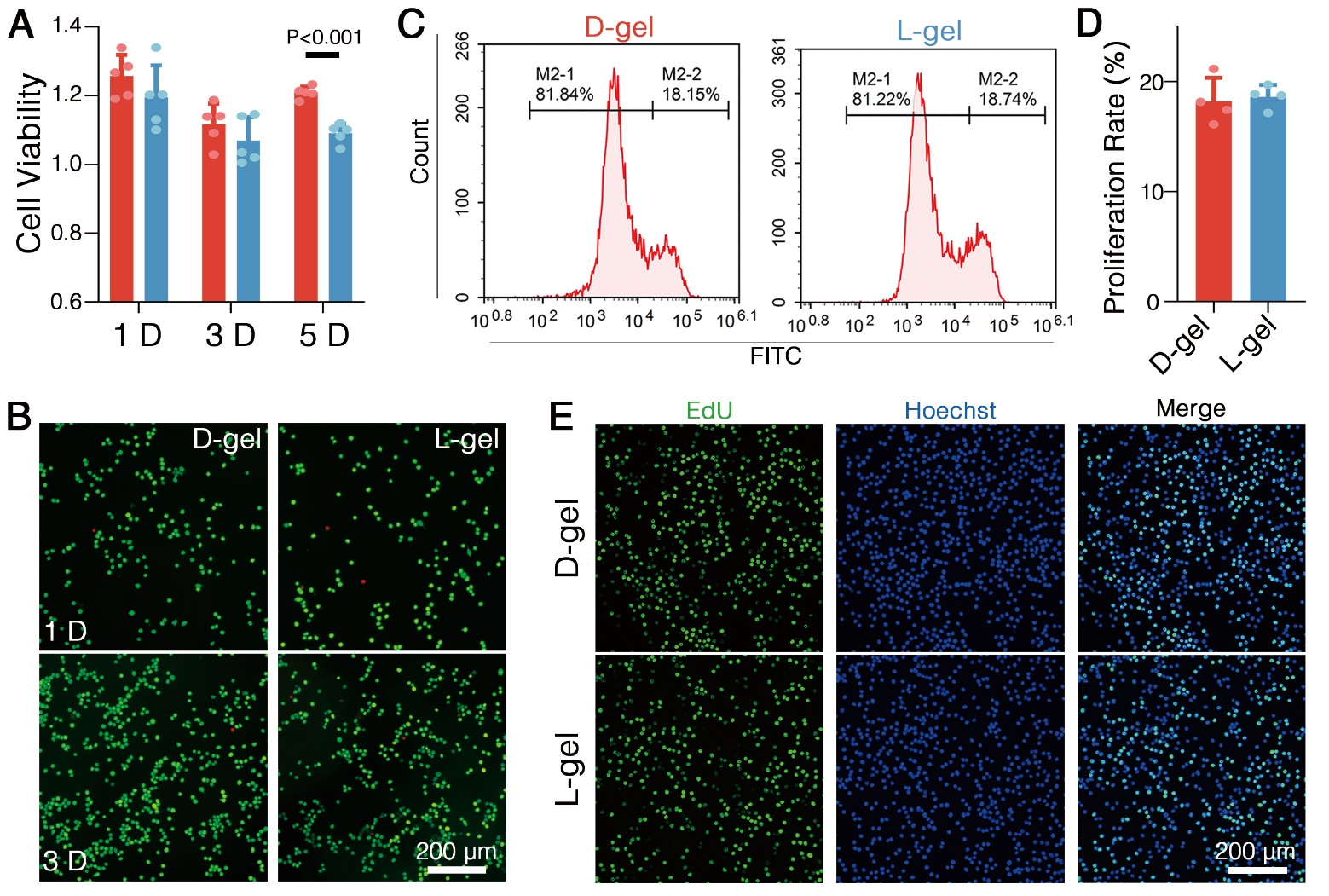

**Fig. S8. Effects of enantiomeric hydrogels on microglia behaviors.** (A) Viability of microglia cultured on enantiomeric hydrogels. (B) Representative micrographics of live/dead staining microglia cultured on enantiomeric hydrogels. (C, D) Representative flow cytometric percentage of EdU-stained proliferating microglia (C) and quantification statistics (D). (E) Representative micrographics of EdU-stained proliferating microglia with hoechst-stained nuclei in blue.

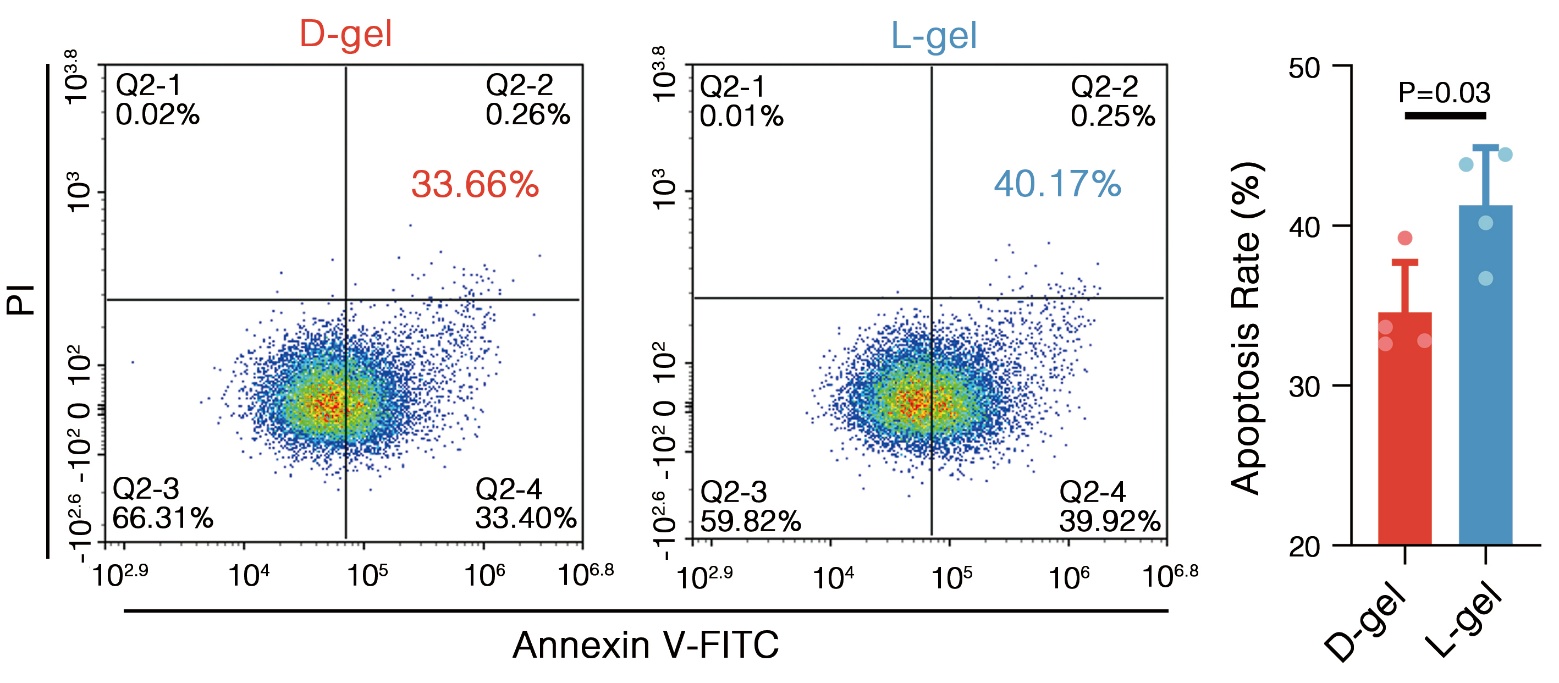

**Fig. S9. Effects of enantiomeric hydrogels on microglia behaviors.** Representative percentage and quantification of apoptotic microglia determined by flow cytometry with Annexin V–FITC/PI staining.

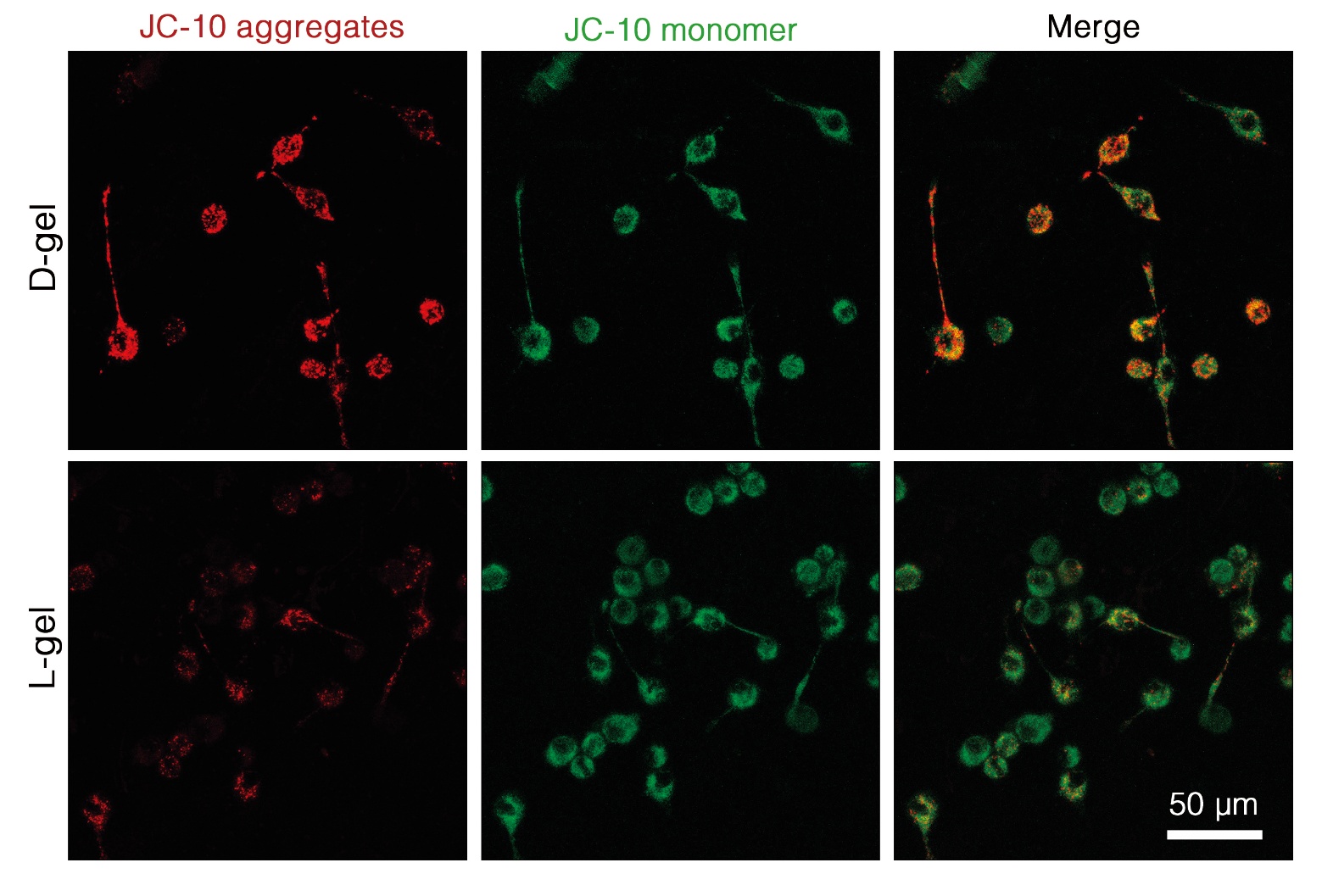

**Fig. S10. Effects of enantiomeric hydrogels on microglia behaviors.** Representative fluorescent micrographics of JC-10-stained microglia, indicating mitochondrial membrane potential.

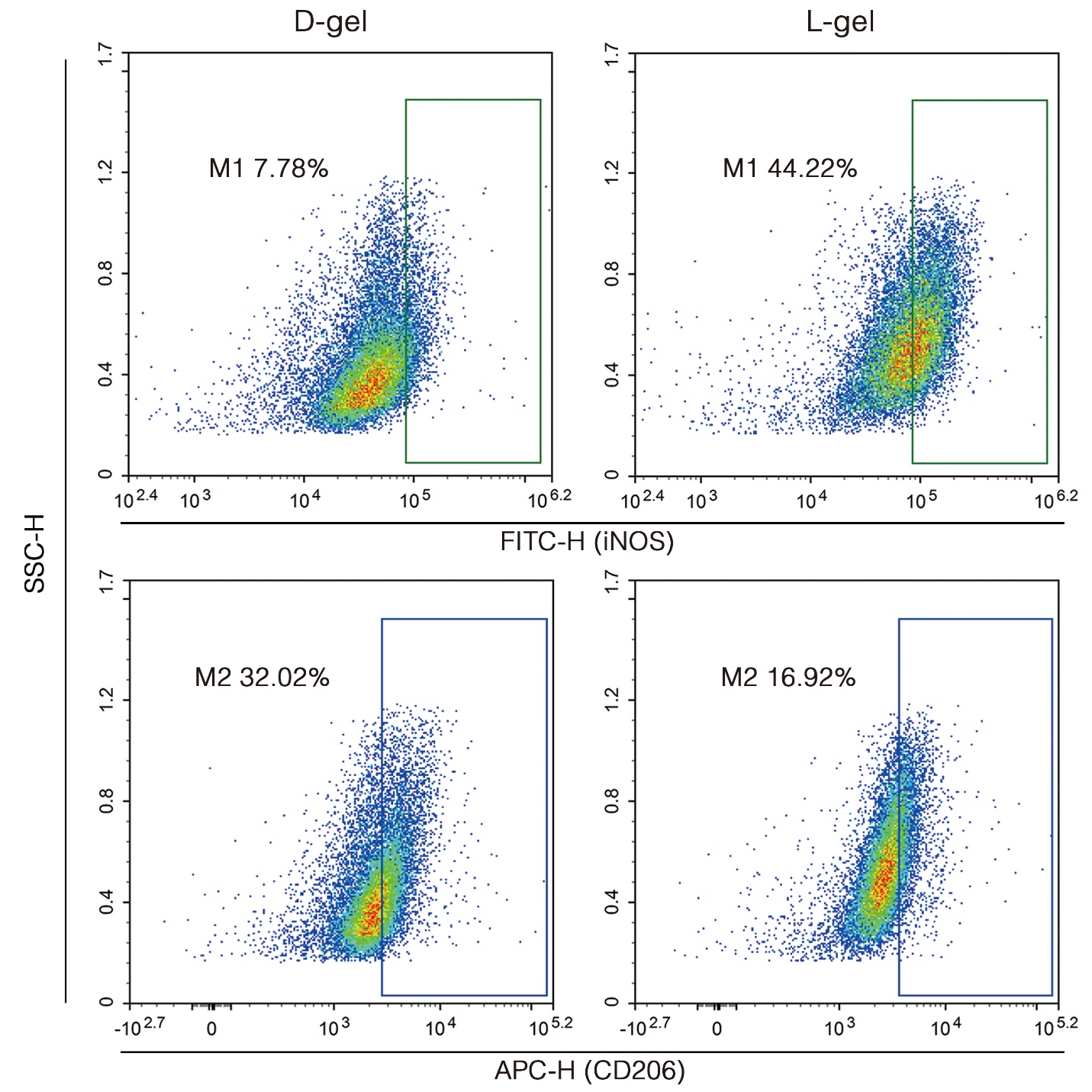

**Fig. S11. Effects of enantiomeric hydrogels on microglial polarization.** Representative percentages of polarized microglia determined by flow cytometry with iNOS and CD206 co-staining. Polarized cells were highlighted in boxes: green indicates M1-polarized cells, and blue indicates M2-polarized cells.

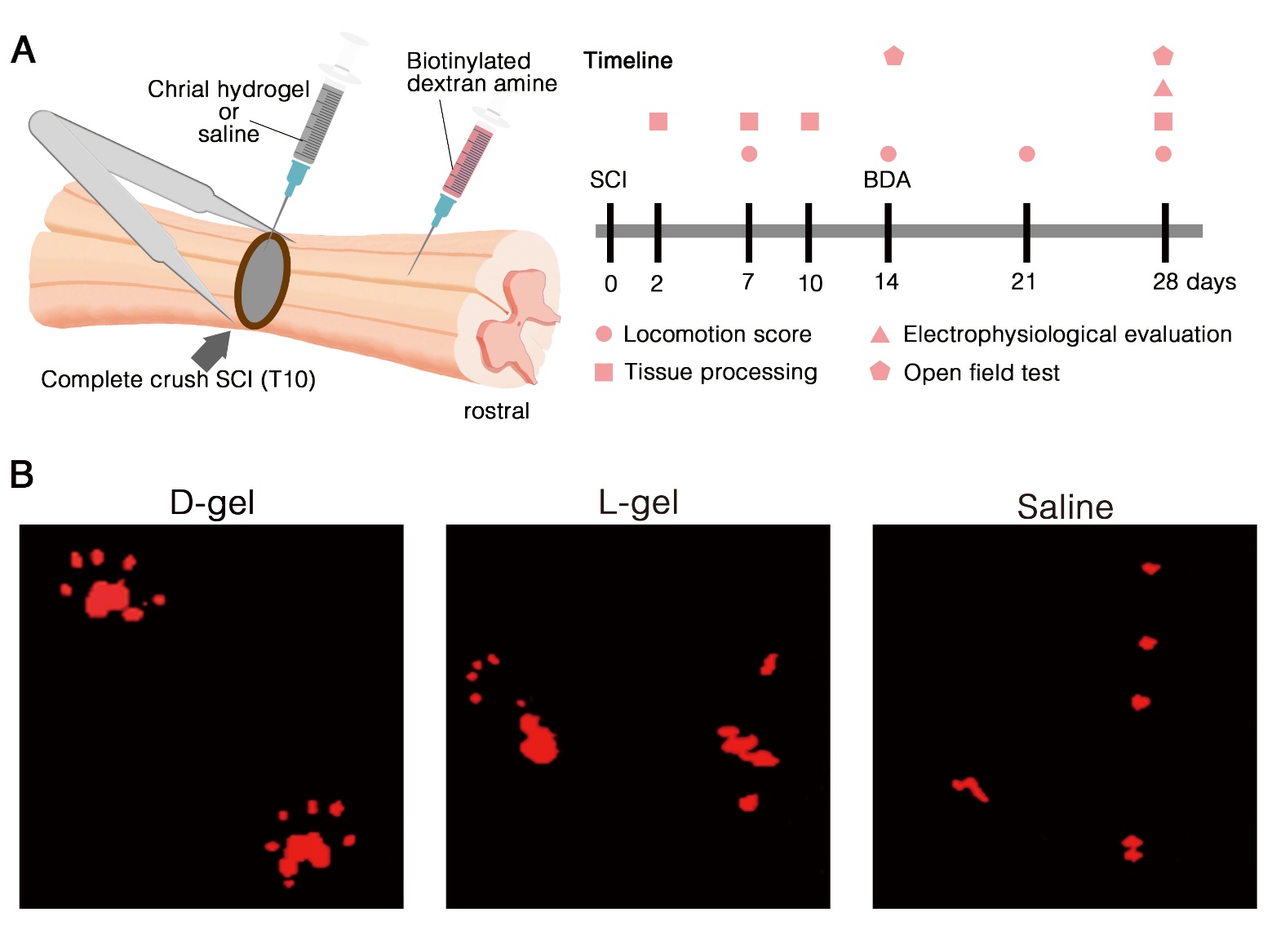

**Fig. S12.** (A) Schematic representation and timeline of animal experiments. (B) Representative locomotor footprints of rats at 28 days.

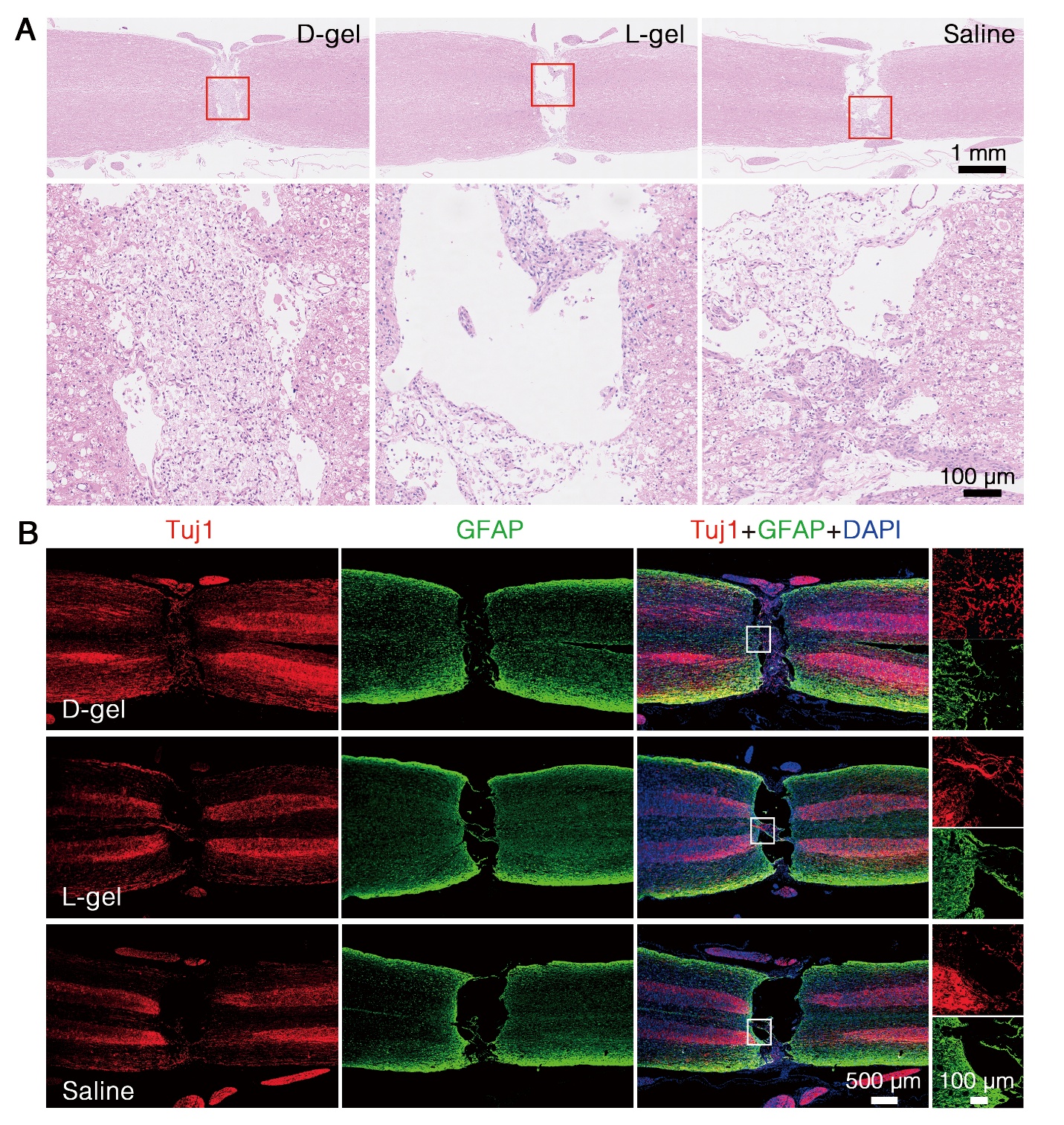

**Fig. S13. Images of injured spinal cords treated with enantiomeric hydrogels.** (A) Photographs of spinal cord lesion at 28 days. (B, C) Representative H&E (B) and IF staining (C) micrographics of longitudinally sectioned spinal cord lesion. Tuj1 labels neurons, GFAP labels astrocytes and DAPI labels nuclei.

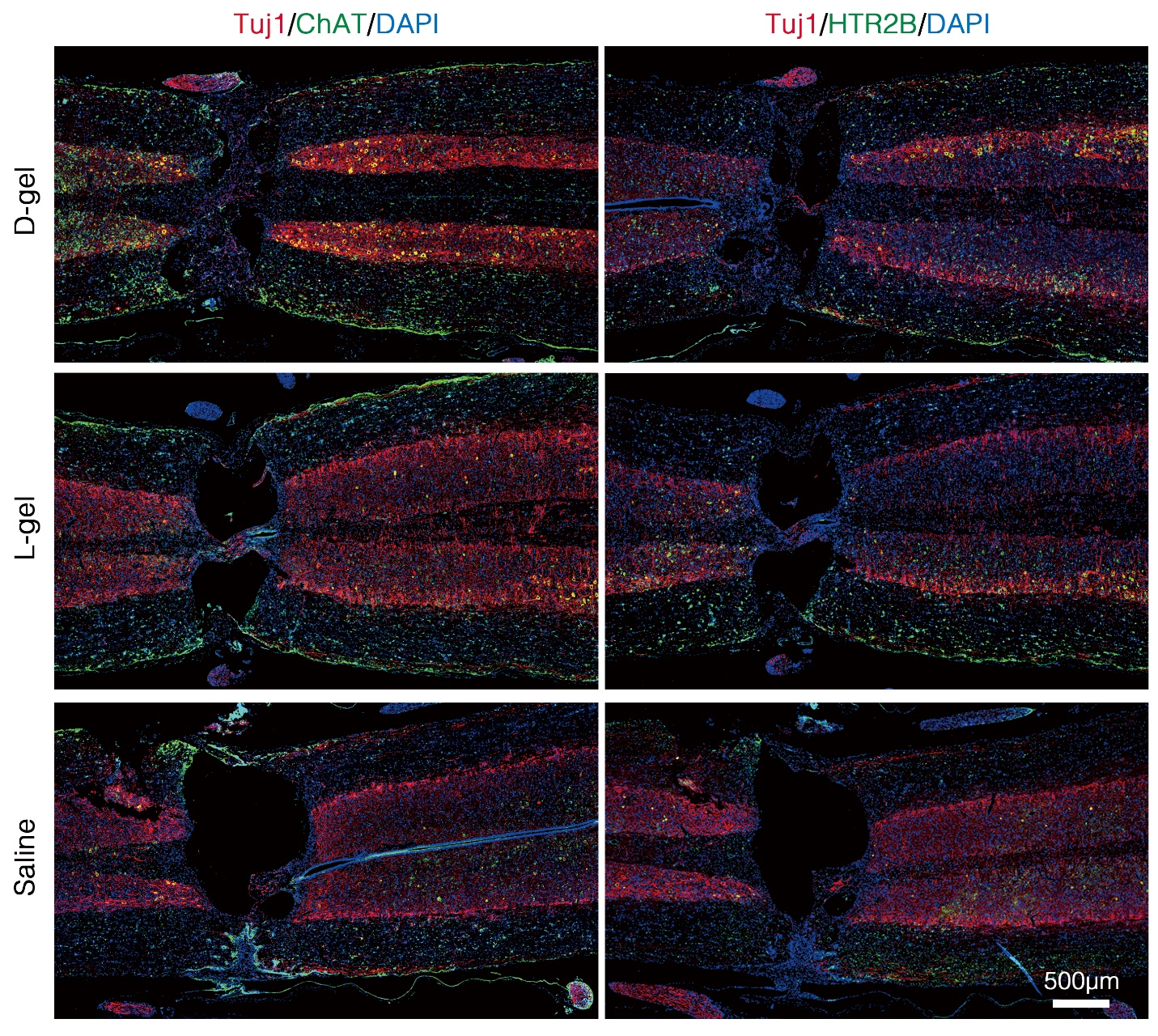

**Fig. S14.** Representative IF staining micrographics of longitudinally sectioned spinal cord lesion for ChAT and HTR2B. Tuj1 labels neurons and DAPI labels nuclei.

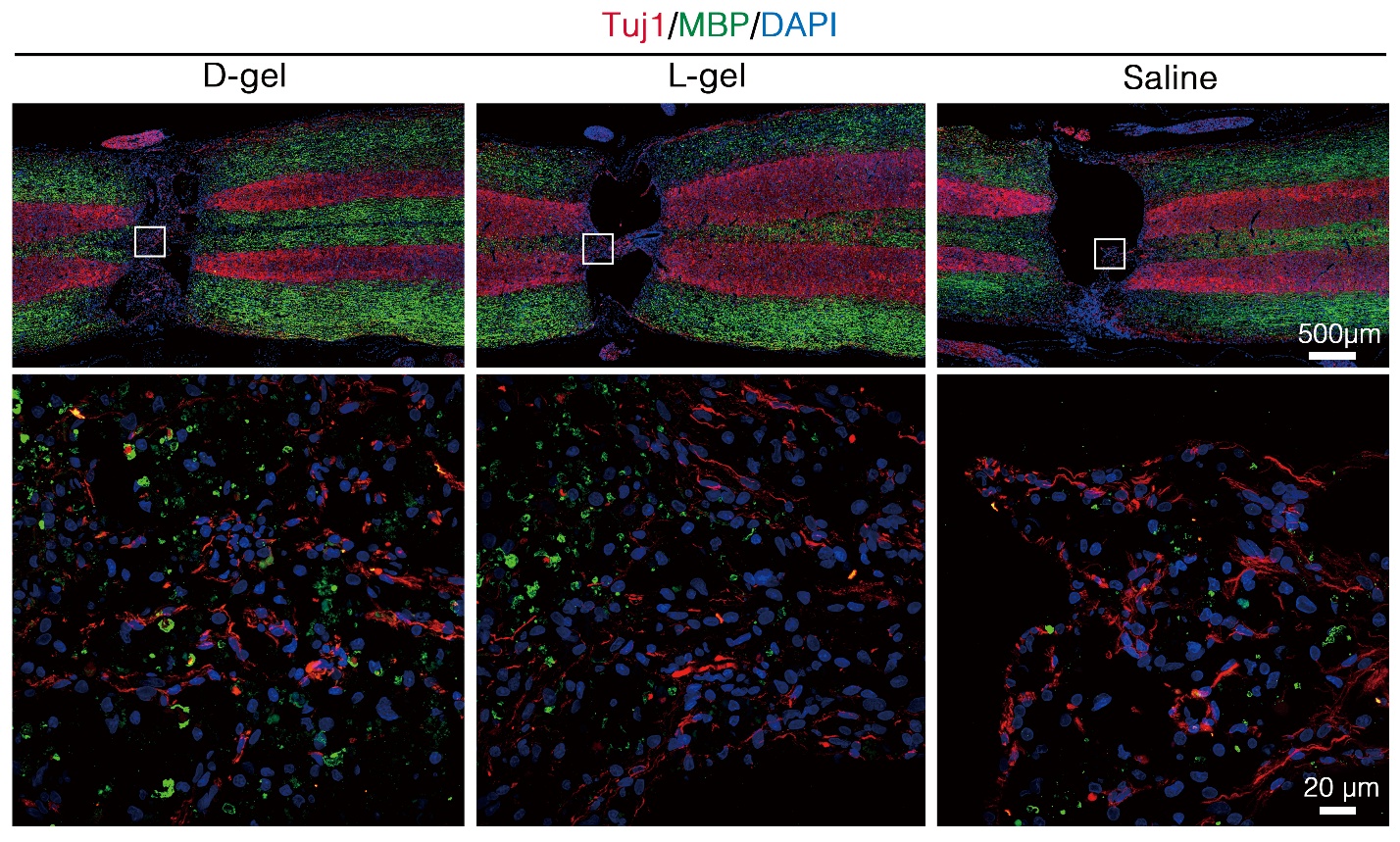

**Fig. S15.** Representative IF staining micrographics of longitudinally sectioned spinal cord lesion for MBP. Tuj1 labels neurons and DAPI labels nuclei.

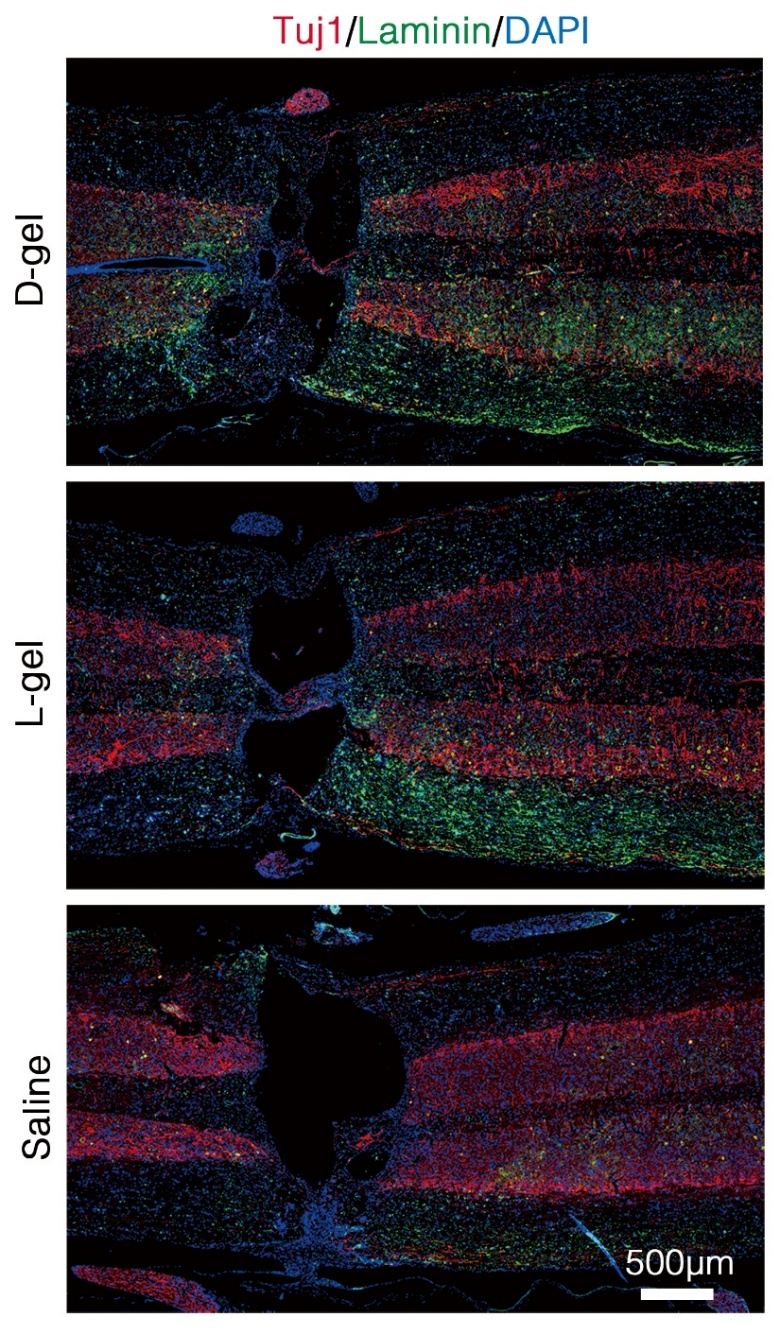

**Fig. S16.** Representative IF staining micrographics of longitudinally sectioned spinal cord lesion for laminin. Tuj1 labels neurons and DAPI labels nuclei.

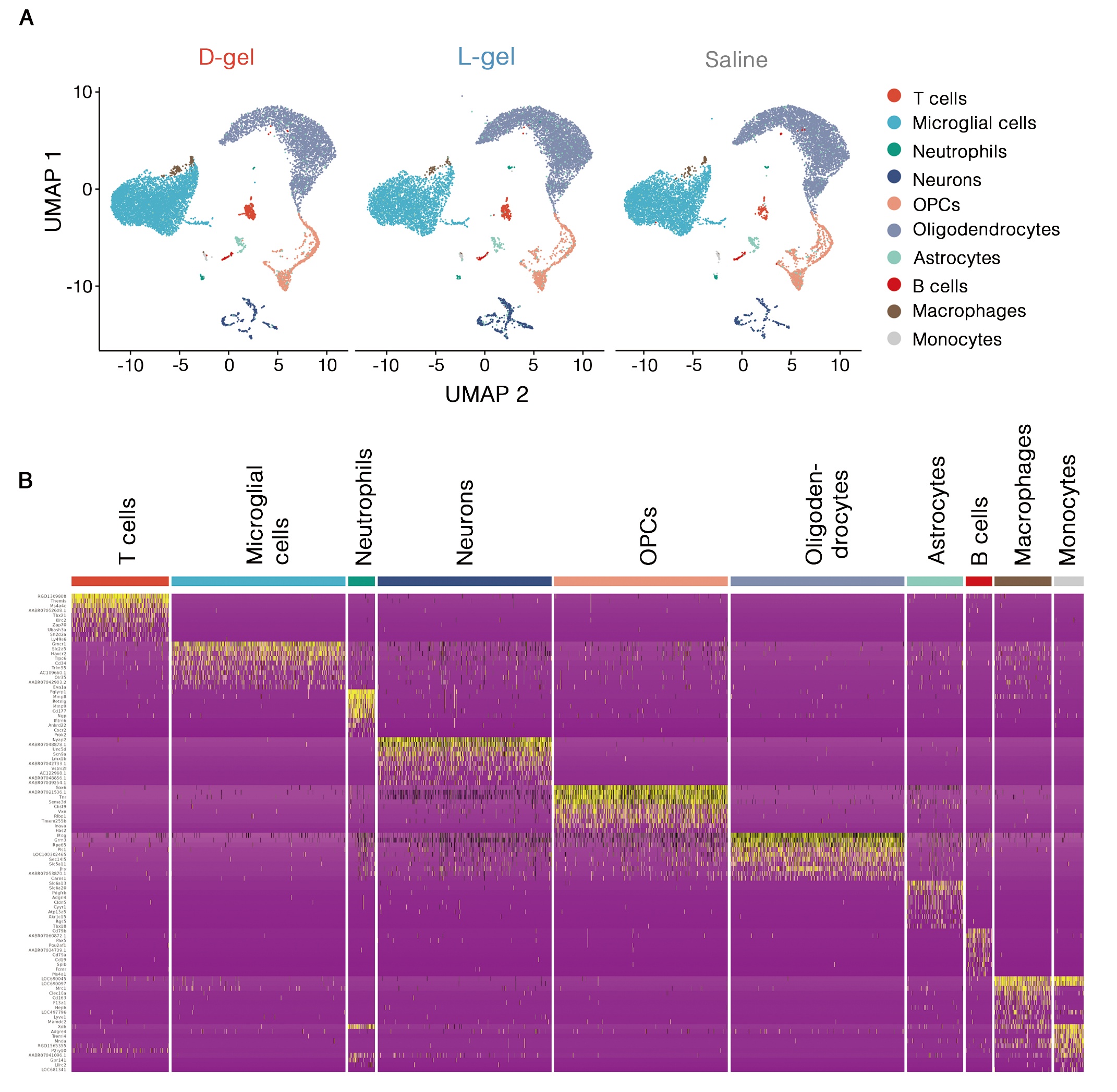

**Fig. S17.** (A) UMAP visualization of ten cell types identified in the injured spinal cord from snRNA-seq atlas. (B) Top 10 marker genes for each cell type.

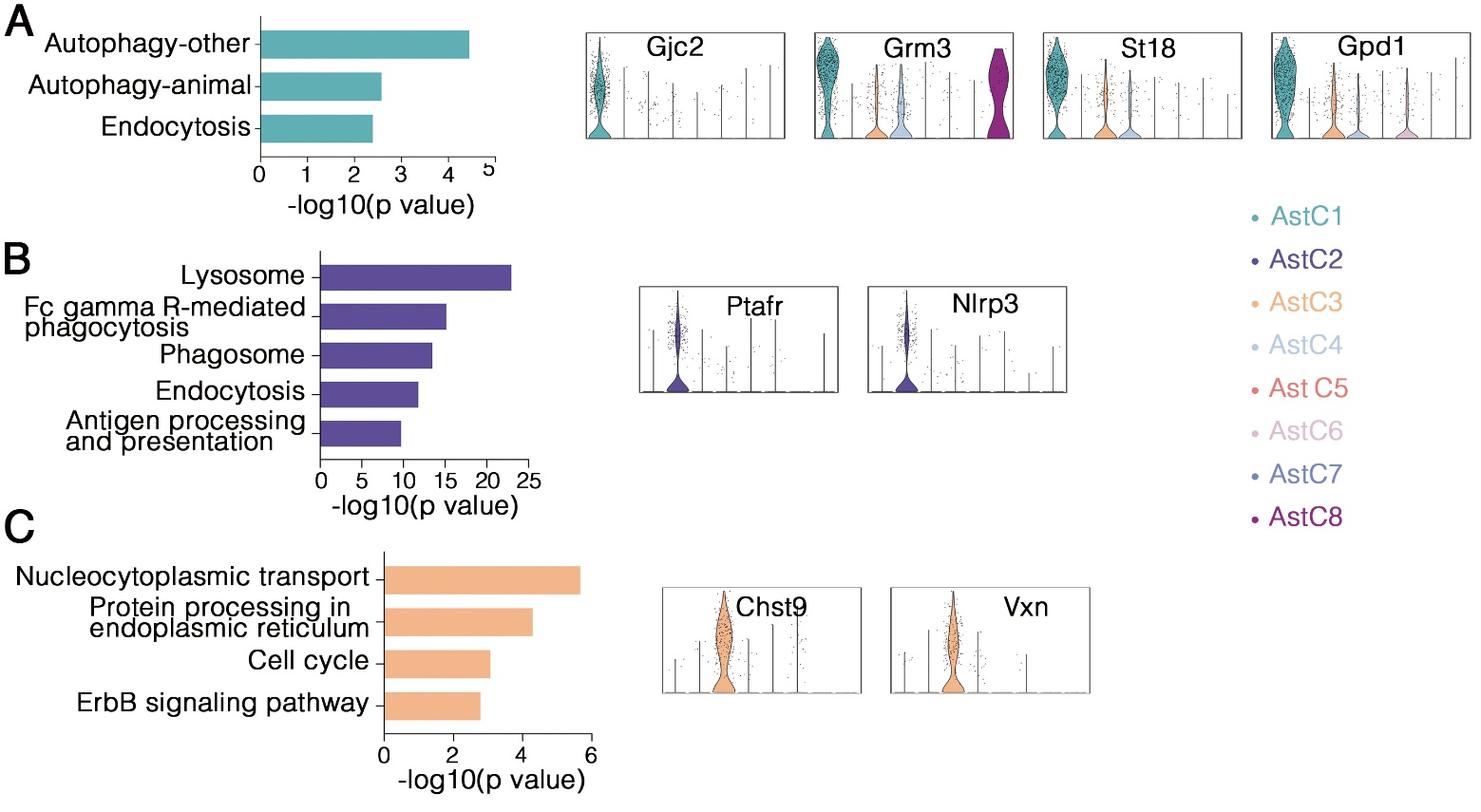

**Fig. S18. Characteristics of astrocyte clusters from snRNA-seq atlas.** Functional annotation of each cluster using enriched KEGG pathways and marker genes. (A) Astrocyte cluster 1. (B) Astrocyte cluster 2. (C) Astrocyte cluster 3.

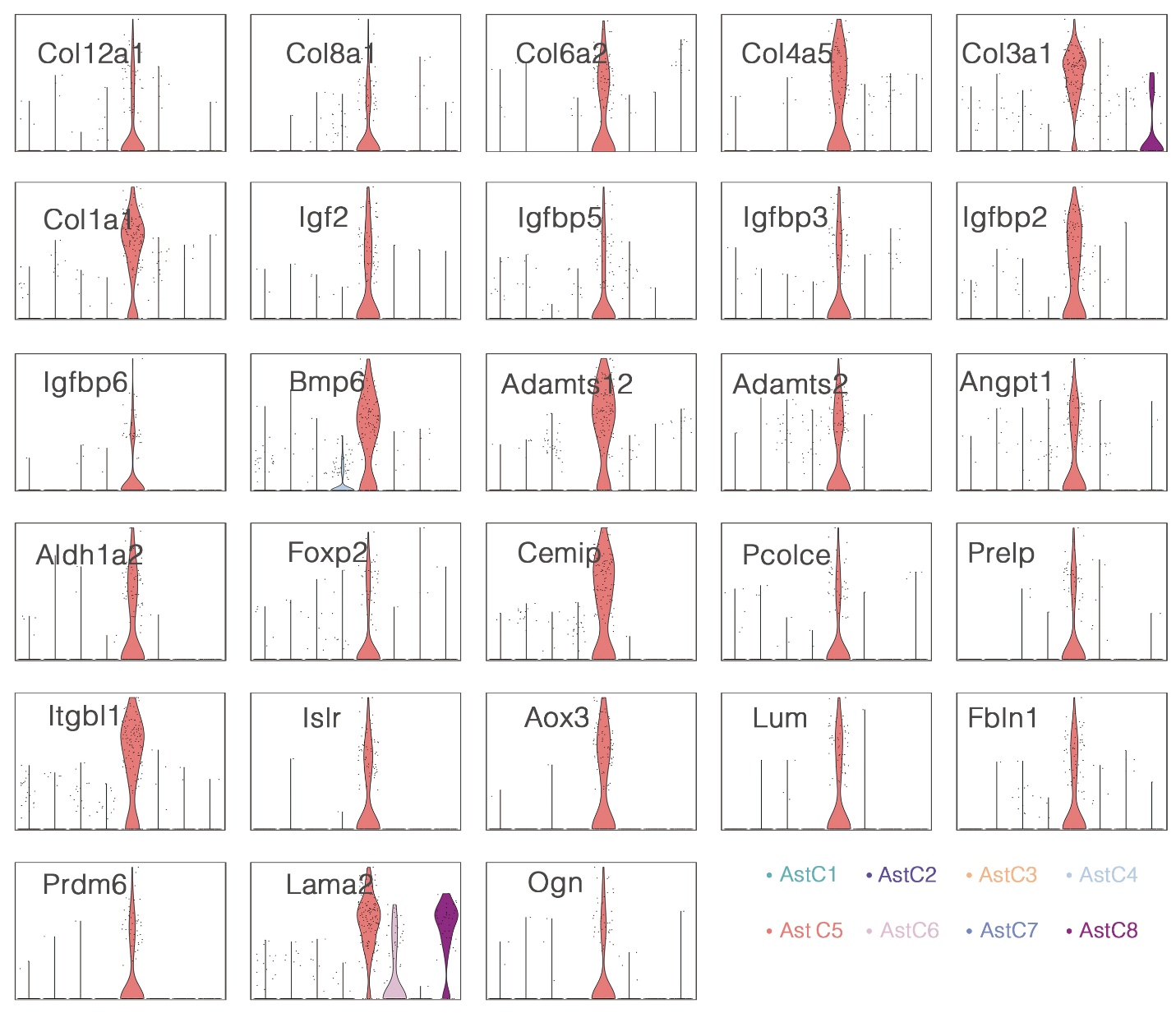

**Fig. S19.** Marker genes of astrocyte cluster 5 from snRNA-seq atlas.

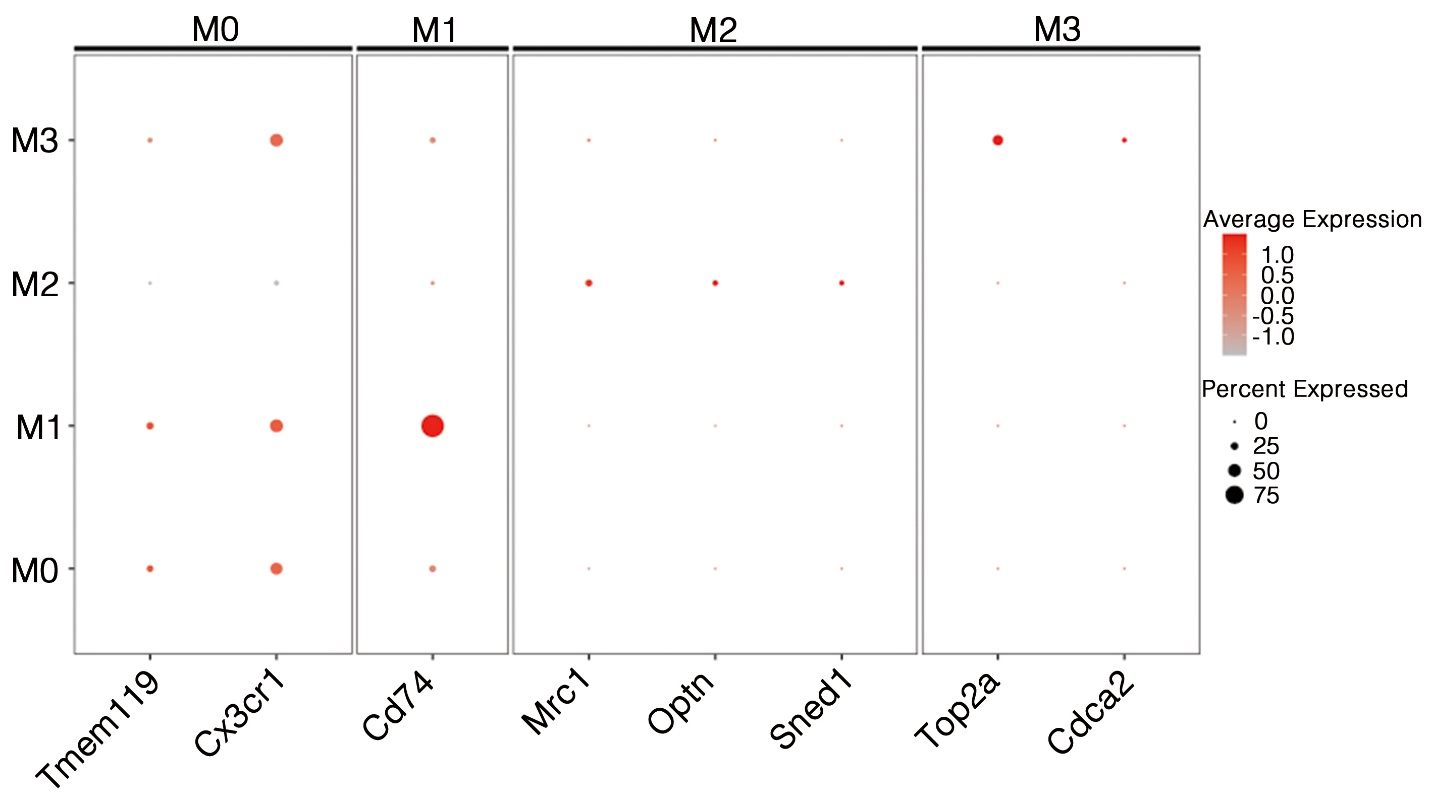

**Fig. S20.** Marker genes used for microglia cluster identification in snRNA-seq atlas.

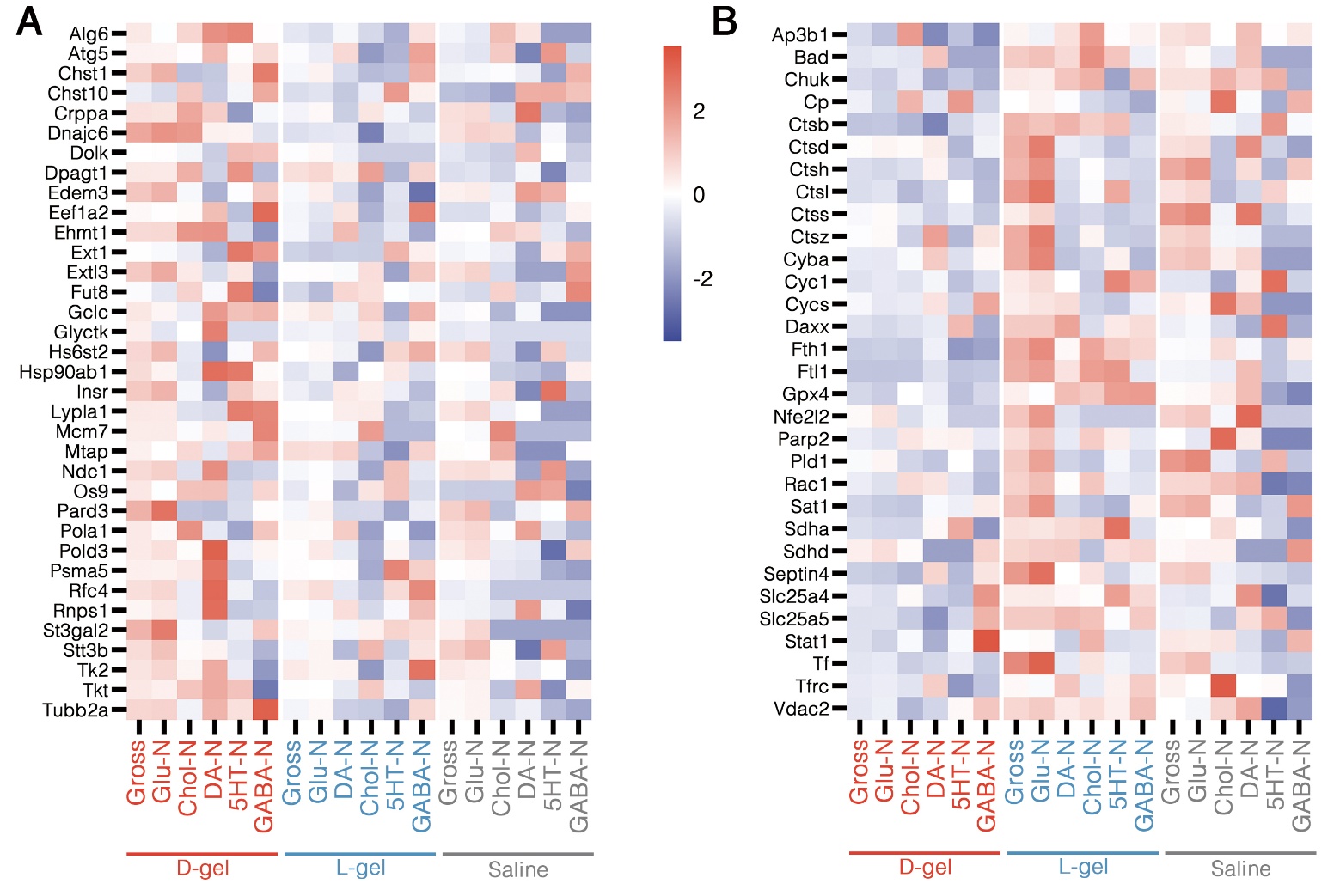

**Fig. S21.** Cellular metabolism and homeostasis-related gene level (A) and cell death and oxidative stress-related gene level (B) heatmaps of neuron clusters and gross across different groups.

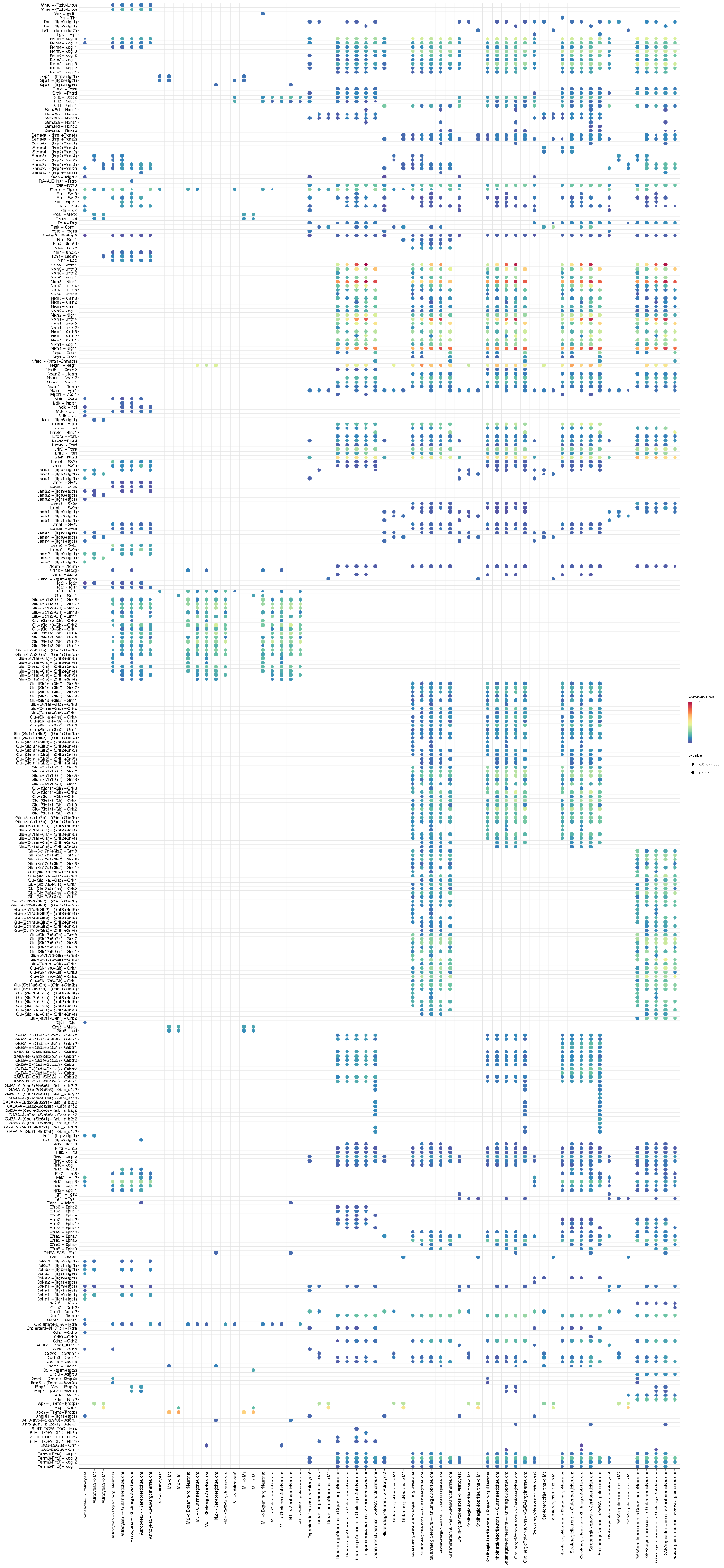

**Fig. S22.** Heatmap of ligand-receptor cell pairs and associated gene pairs in the D-gel group.

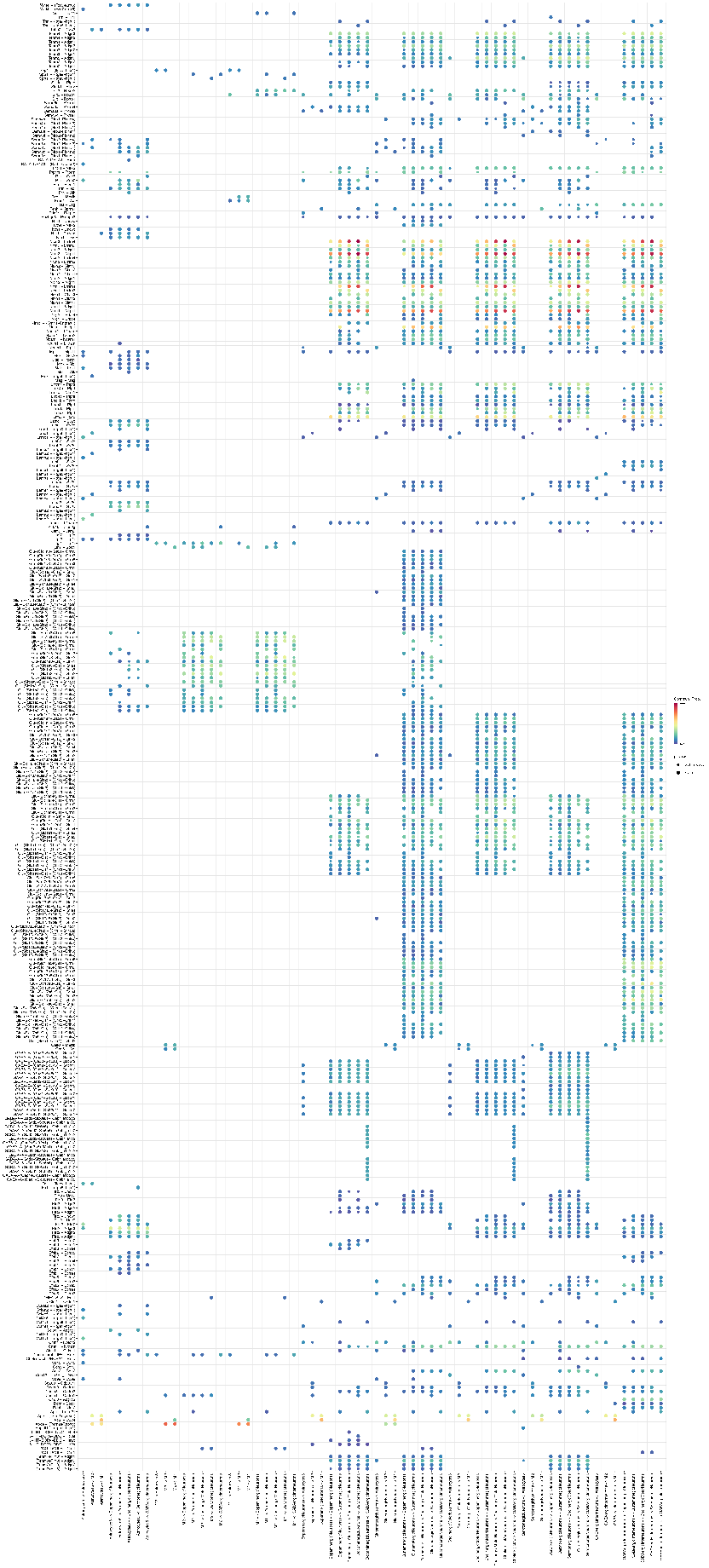

**Fig. S23.** Heatmap of ligand-receptor cell pairs and associated gene pairs in the L-gel group.

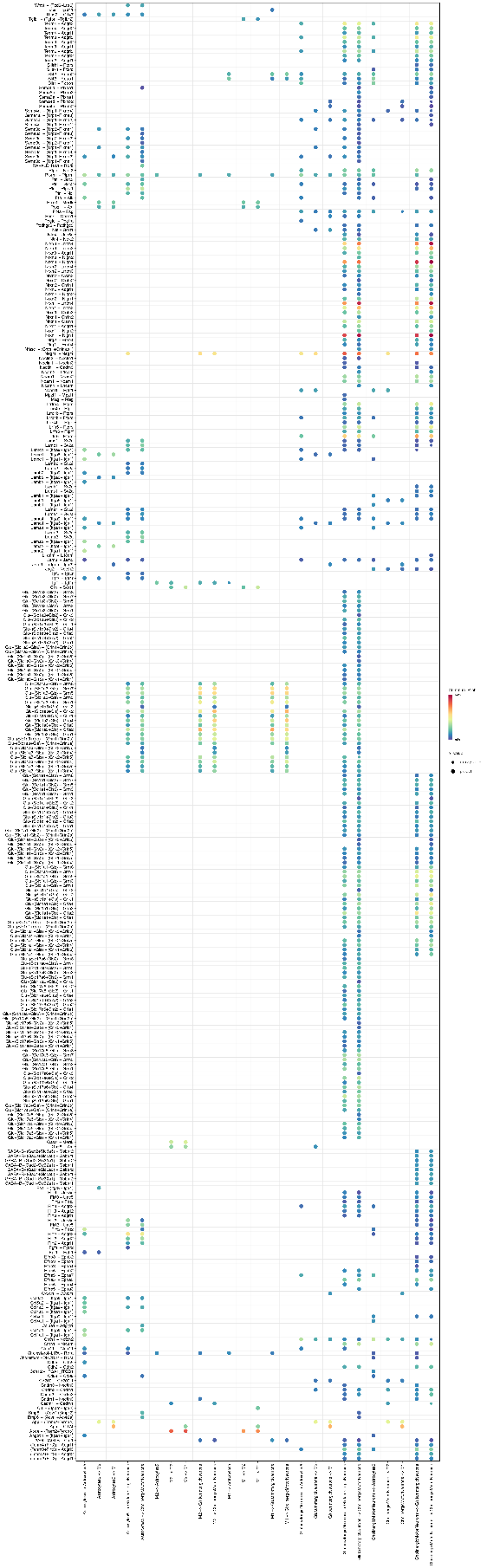

**Fig. S24.** Heatmap of ligand-receptor cell pairs and associated gene pairs in the saline group.

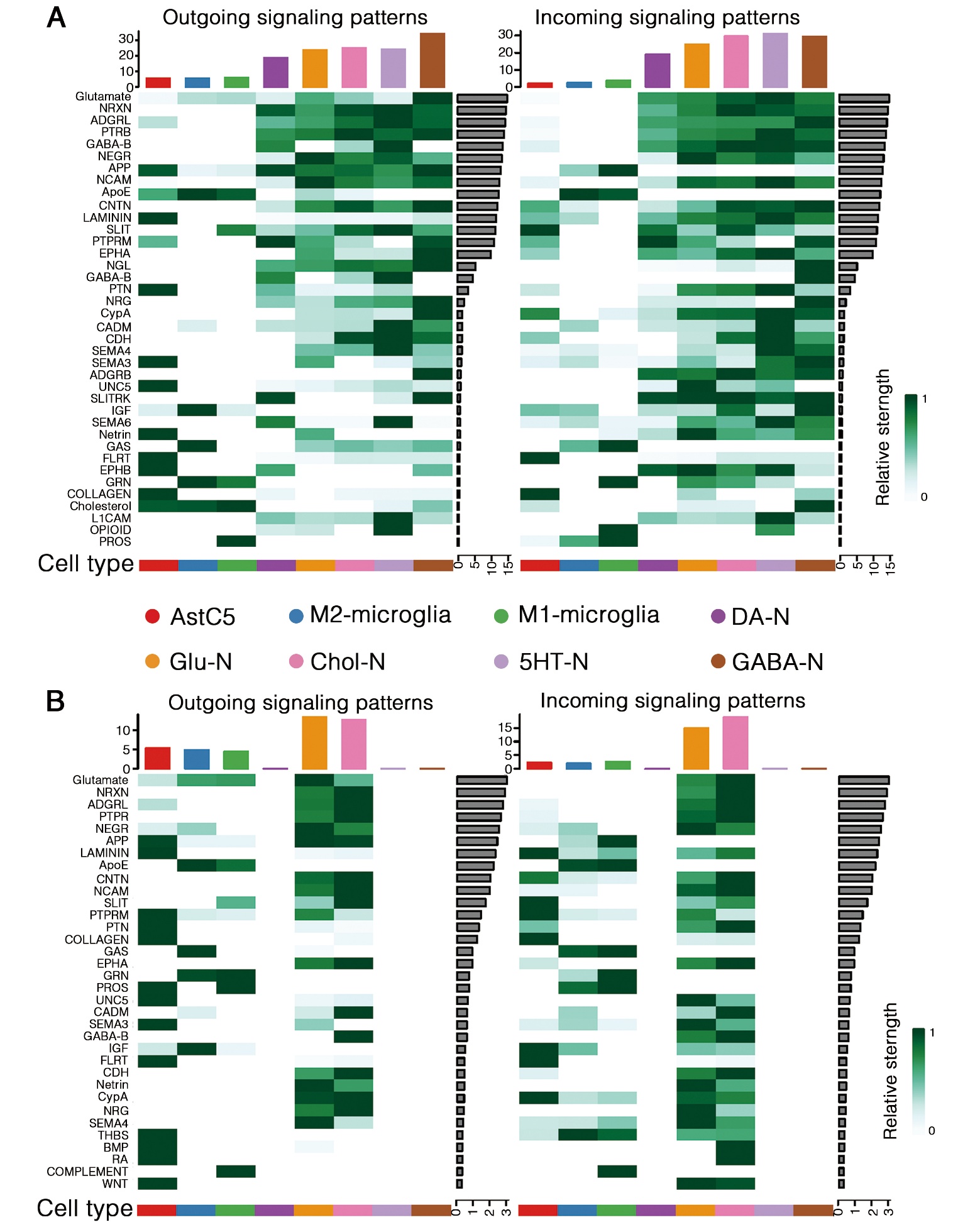

**Fig. S25.** High contributed signaling pathway patterns of the L-gel group (A) and saline group (B) in cellular communication. Outgoing signaling patterns indicate ligand-providing cell types, and incoming signaling patterns indicate receptor-providing cell types.

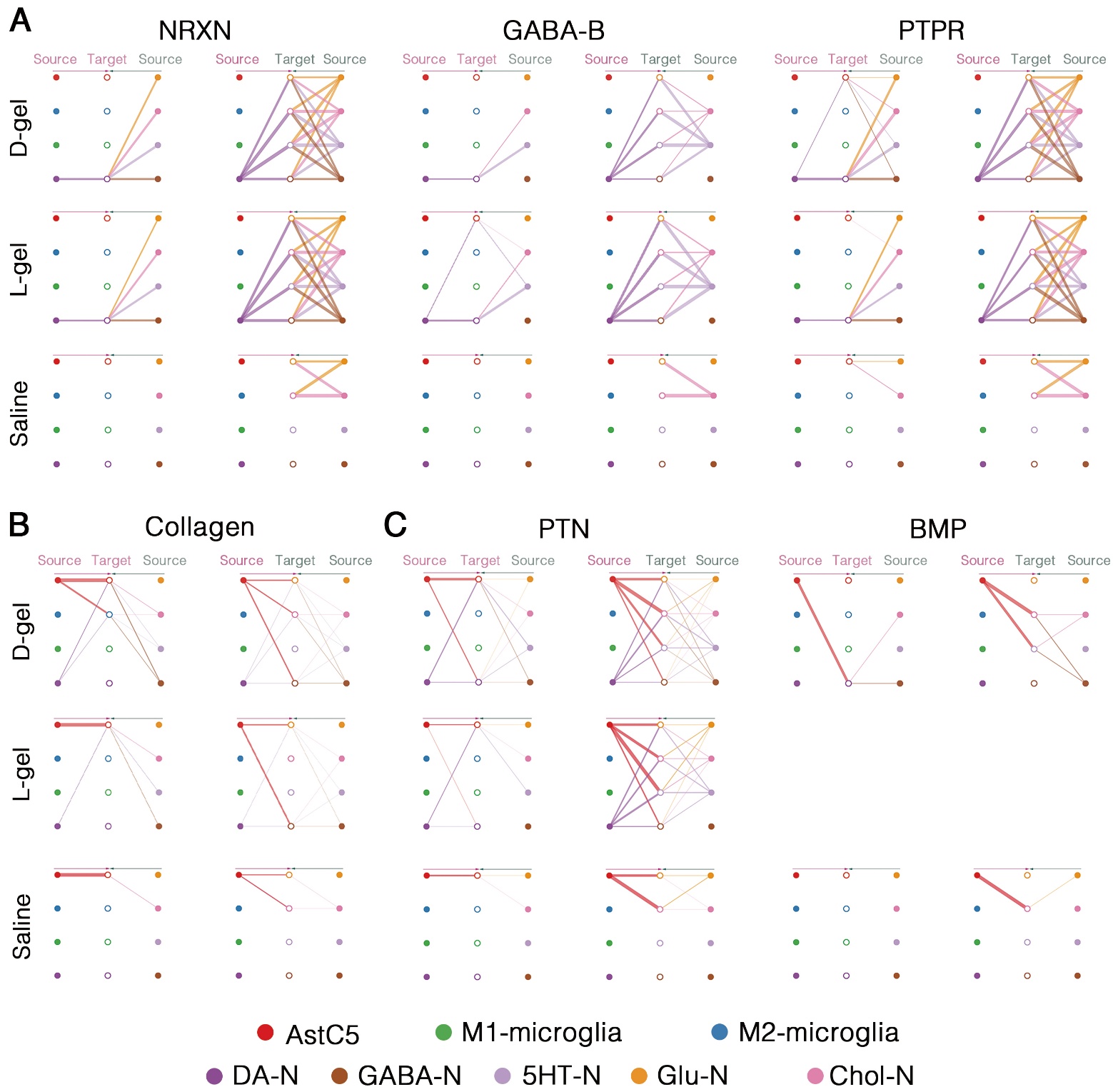

**Fig. S26. Hierarchical diagram of ligand-receptor interactions within signaling pathways:** pathways between neurons (A), pathways through which AstC5 affects others (B), pathways through which AstC5 influences neurons exclusively (C). Solid circles represent ligand-providing cells, hollow circles represent receptor-providing cells, and thicker lines indicate stronger signals.

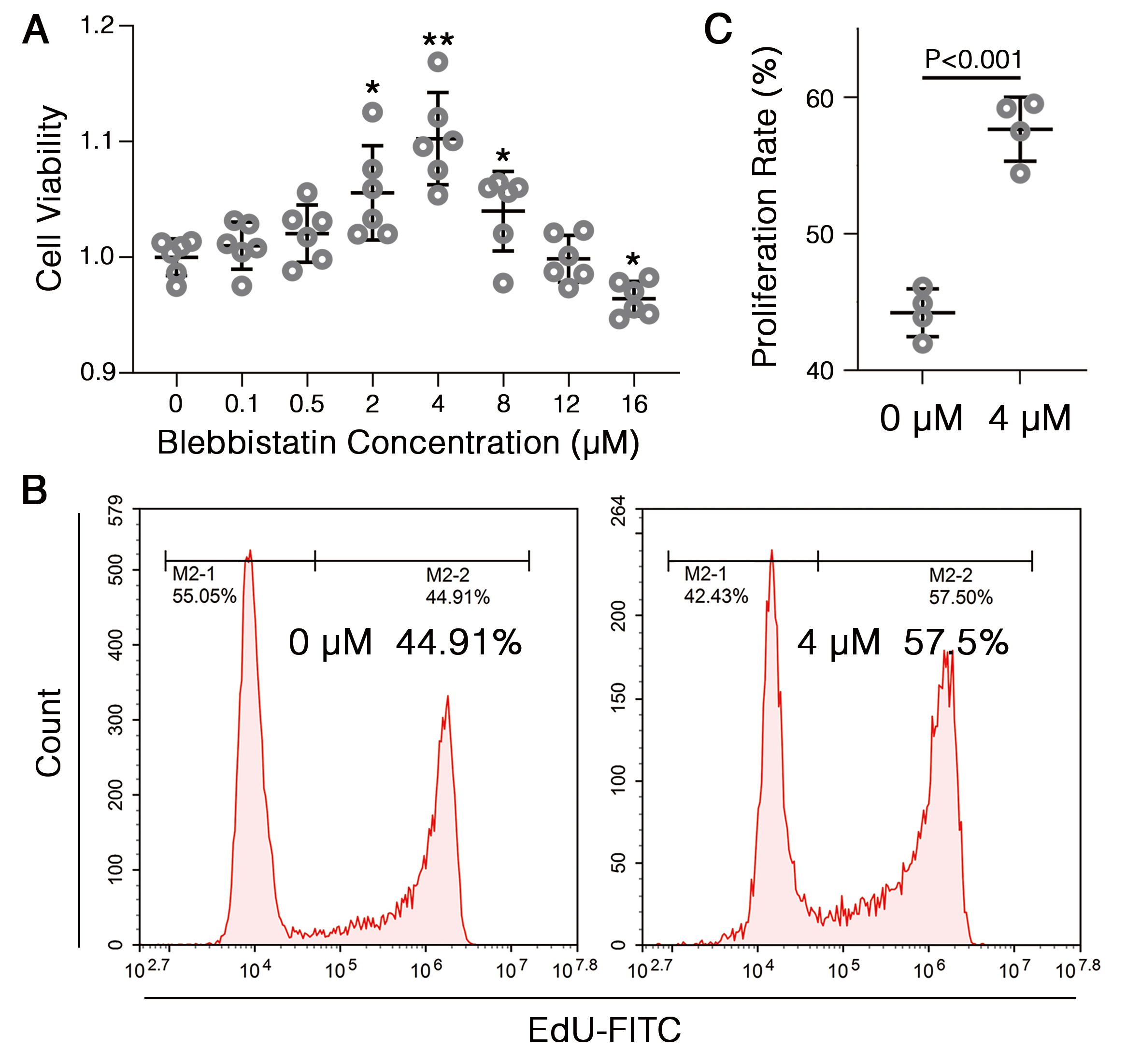

**Fig. S27.** (A) Cell viability of astrocytes cultured at different blebbistatin concentration, **P<0.05*, ***P<0.01*, compared with 0 μM group. (B, C) Representative flow cytometric percentage of EdU-stained proliferating astrocytes (B) and quantification statistics (C) at 0 and 4μM blebbistatin.

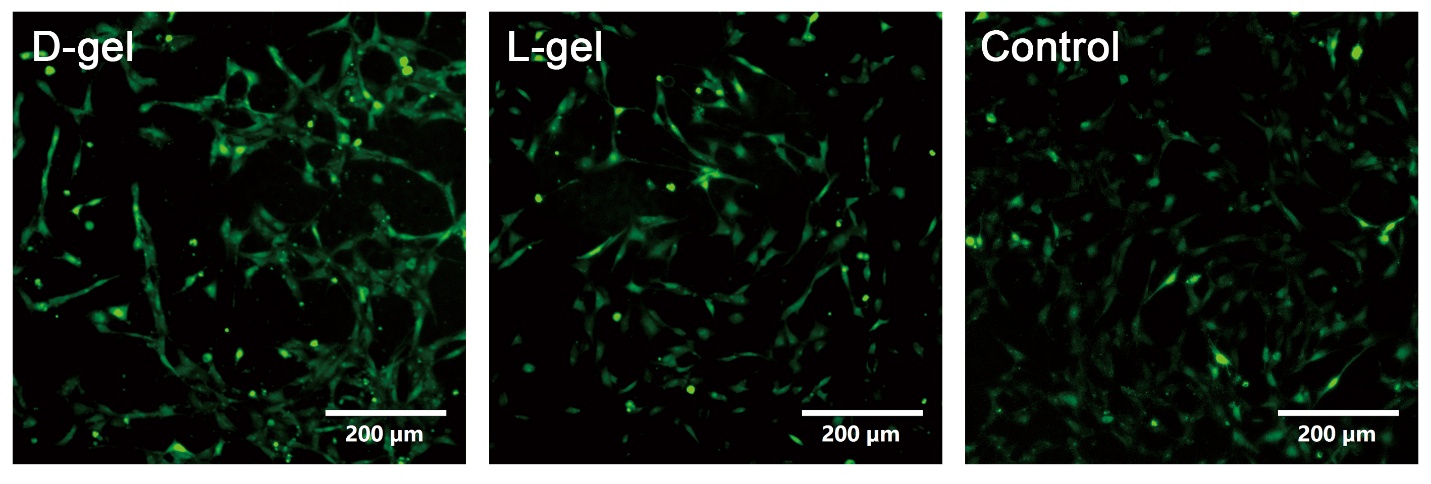

**Fig. S28.** Representative fluorescence imaging of stained Ca^2+^ in astrocytes.
