## Supplementary figures and images for "Enantiomeric Hydrogel-Manipulated Mechanotransduction Triggers Neurogenesis and Immunomodulation for Spinal Cord Repair"

### Supplemental figure 22

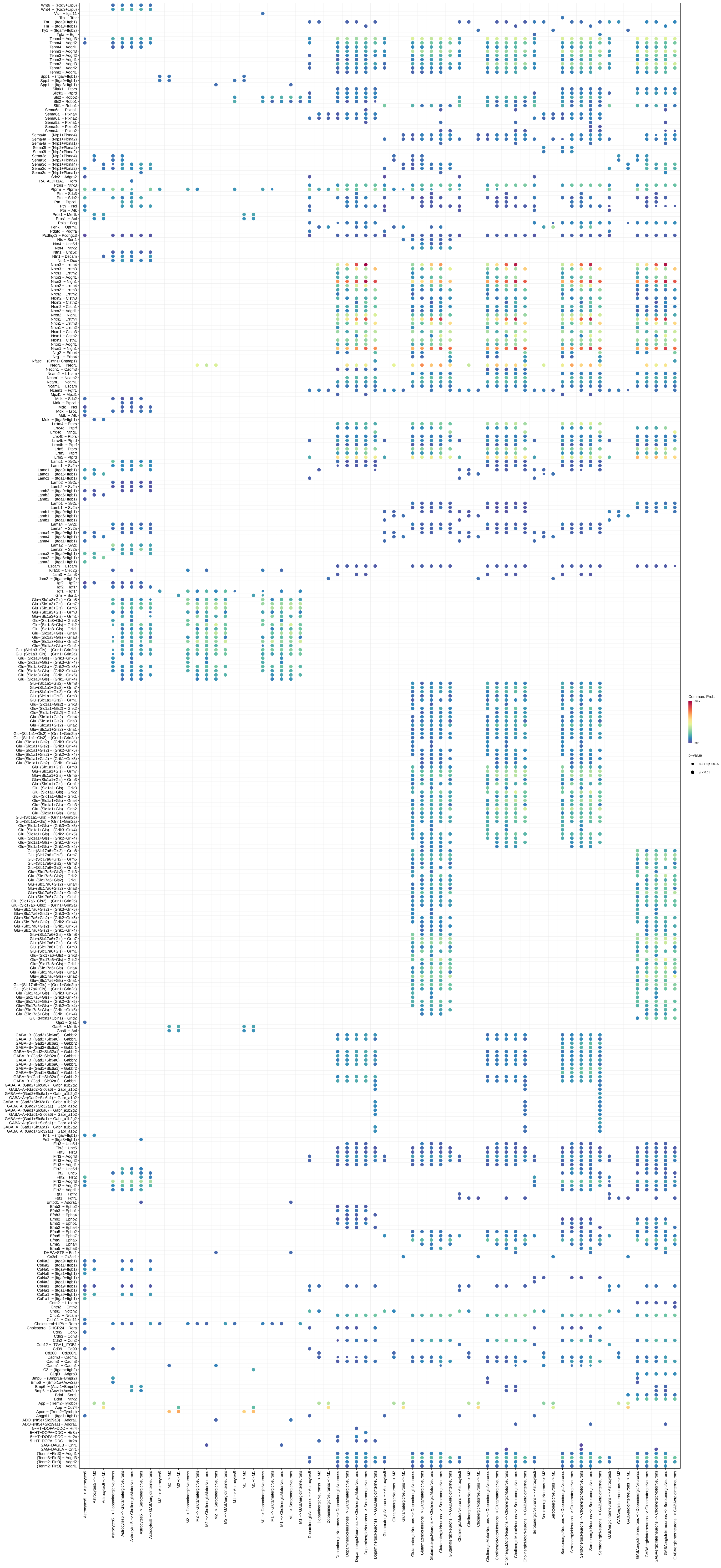

### Supplemental figure 23

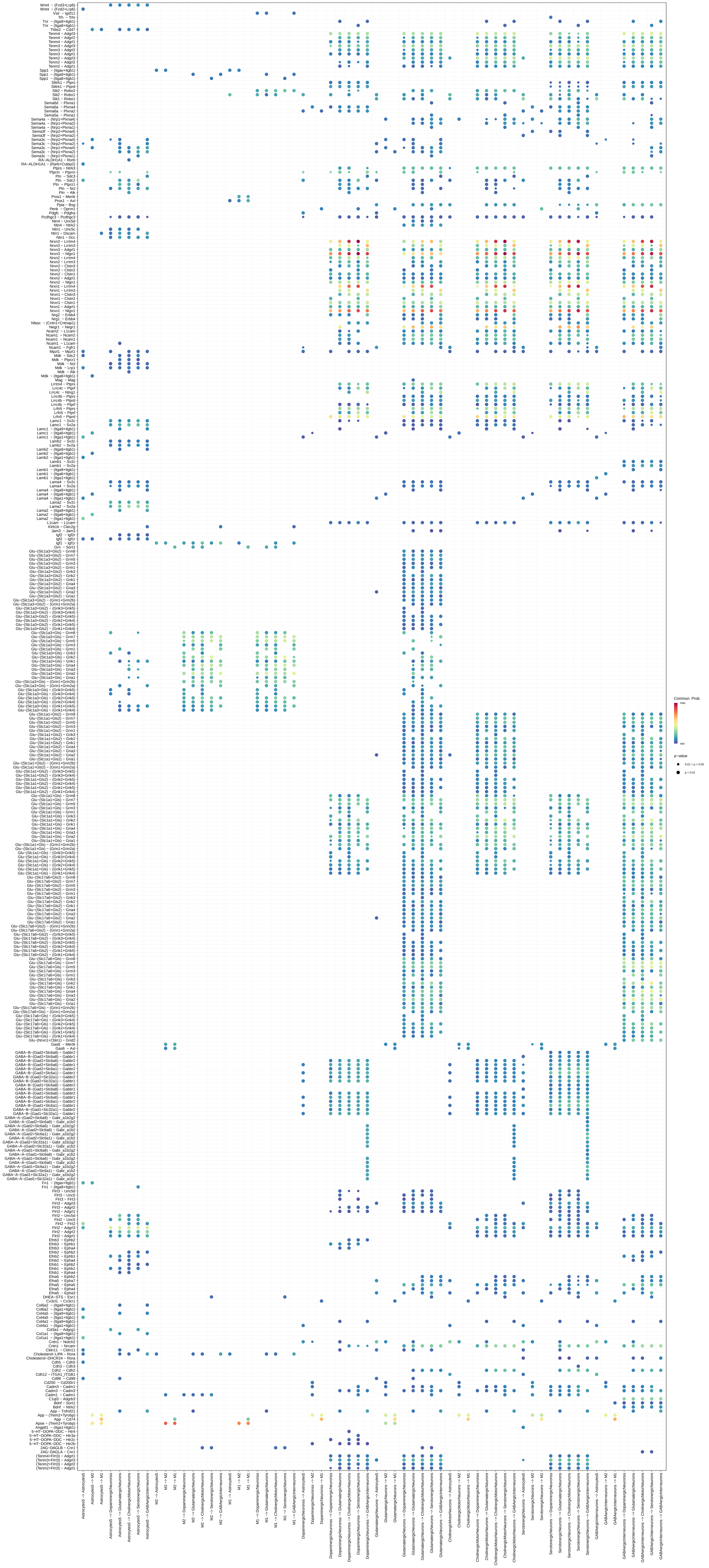

### Supplemental figure 24

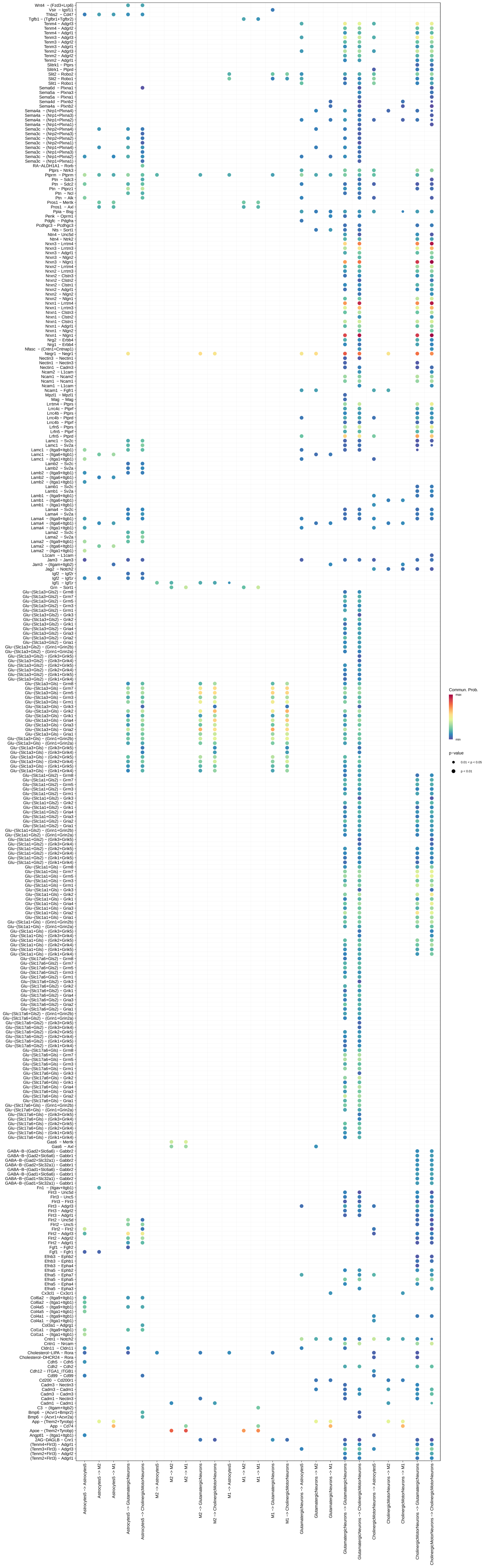
